# Mammalian cells internalize bacteriophages and utilize them as a food source to enhance cellular growth and survival

**DOI:** 10.1101/2023.03.10.532157

**Authors:** Marion C. Bichet, Jack Adderley, Laura Avellaneda, Linden J. Gearing, Celine Deffrasnes, Cassandra David, Genevieve Pepin, Michael P. Gantier, Ruby CY Lin, Ruzeen Patwa, Gregory W. Moseley, Christian Doerig, Jeremy J. Barr

## Abstract

There is a growing appreciation that the direct interaction between bacteriophages and the mammalian host can facilitate diverse and unexplored symbioses. Yet the impact these bacteriophages may have on mammalian cellular and immunological processes is poorly understood. Here we applied highly purified phage T4, free from bacterial by-products and endotoxins to mammalian cells and analyzed the cellular responses using luciferase reporter and antibody microarray assays. Phage preparations were applied *in vitro* to either A549 lung epithelial cells, MDCK-I kidney cells, or primary mouse bone marrow derived macrophages with the phage-free supernatant serving as a comparative control. Highly purified T4 phages were rapidly internalized by mammalian cells and accumulated within macropinosomes but did not activate the inflammatory DNA response TLR9 or cGAS-STING pathways. Following eight hours of incubation with T4 phage, whole cell lysates were analyzed via antibody microarray that detected expression and phosphorylation levels of human signaling proteins. T4 phage internalization led to the activation of AKT-dependent pathways, resulting in an increase in cell metabolism, survival, and actin reorganization, the last being critical for macropinocytosis and potentially regulating a positive feedback loop to drive further phage internalization. T4 phages additionally down-regulated CDK1 and its downstream effectors, leading to an inhibition of cell cycle progression and an increase in cellular growth through a prolonged G1 phase. These interactions demonstrate that highly purified T4 phages do not activate DNA-mediated inflammatory pathways but do trigger protein phosphorylation cascades that promote cellular growth and survival. We conclude that mammalian cells are internalizing bacteriophages as a food source to promote cellular growth and metabolism.

## INTRODUCTION

Bacteriophages, also called phages, are viruses that infect and kill bacteria, their natural hosts. Phages are ubiquitous across environments and are intrinsic components of our microbiomes, colonizing all niches of the body (1). As such, the human body is frequently and continuously exposed to a diverse community of phages(2,3). This is especially true within the gut, which houses a high-diversity microbial community(4). Phages are essential components of the gut and participate in the genetic diversification and individualization of the gut microbiome throughout our lifespan (5–10). While phages facilitate many changes to these gut microbial communities, they are also known to interact with the underlying mammalian cells (1,11–14). Mammalian cells can engulf phages via a variety of mechanisms, leading to the internalization and accumulation of active phages (12–15). Phages have been shown to bind specific mammalian cellular receptors, triggering receptor-mediated endocytosis (16,17). However, this mechanism appears to be quite rare, as the probability of having matching receptors and ligands between mammalian cells and phages is low. The predominant mechanism by which phages have been shown to enter mammalian cells is non-specific internalization via macropinocytosis (1, 13, 14).

Macropinocytosis is an actin-based process that is a unique pathway of endocytosis characterized by the non-specific internalization of extracellular fluid, nutrients, and potential microorganisms in large endocytic vesicles (50–1000 nm) known as macropinosomes. Macropinocytosis is essential for cellular growth and cell proliferation as it allows the cell to gain large amounts of extracellular nutrients (18–22). Additionally, macropinocytosis can be utilized to sample and subsequently detect pathogens and foreign nucleic acids, leading to the activation of the innate immune system. This immune sensing of foreign nucleic acids is a hallmark of antiviral defense; it is a highly regulated process that involves cellular compartmentalization and the selective recognition of foreign nucleic acids (23). Macropinocytosis of phages by mammalian cells is a non-specific process whereby cells create actin-mediated ruffles elongating from the cytosol toward the environment, which engulf the extracellular milieu and any phages residing within it. Phages internalized via this pathway steadily accumulate within intracellular macropinosomes (14). The downstream processing of the macropinosome can follow various pathways, including fusion with other endocytic vesicles, fusion with lysosomes leading to acidification and the inactivation of internalized components, recycling and transport to plasma membranes, and constitutive exocytosis (24).

Once inside the cell, phages may stimulate a diverse array of effects. The few studies that have investigated the cellular and innate immune response to phages have hinted at two opposing responses. On the one hand, certain phages are known to induce anti-inflammatory effects (24,25). This has led to suggestions that phages could be used in transplantation patients to reduce risks of organ rejection (25,26) and may play an immunomodulatory role in the gut microbiome (27,28). On the other hand, a growing number of studies have shown pro-inflammatory immune responses and inflammation in response to specific phages (29–31). Collectively, these studies demonstrate that certain phages can induce anti- or pro-inflammatory innate immune responses and highlight an underlying specificity for the cellular detection of specific viral types. Despite growing evidence in the field, it remains mechanistically unclear how phages interact with and modulate the mammalian cells’ innate immune response, and how these interactions can influence downstream cellular processes. Conceptually, the fundamental question as to why mammalian cells are internalizing phage particles and what selective advantage this pertains, remains open.

In this study, we investigate whether phage T4 can modulate the cellular and innate immune pathways across two cell lines *in vitro*. We demonstrate that phages were internalized by mammalian cells via macropinocytosis, with functional phages continually accumulating within macropinosomes ((13,14). All phage preparations were highly purified and confirmed to be free of bacterial endotoxins (32,33). To further ensure the cellular responses detected were elicited by the phages themselves, we used an additional comparative control that consisted of the highly purified phage lysate filtered four times through a 0.02 μm filter to remove phage particles and obtain a phage-free lysate composed of the background supernatant. Using these samples, we then performed luciferase assays and interrogated antibody microarrays to probe the cellular and innate immune changes induced by the presence of T4 phages.

## RESULTS

### T4 phage does not activate the intracellular DNA-sensing receptors TLR9 and cGAS-STING

We focused on bacteriophage T4, a virulent *Tevenvirinae* phage ~200 nm long with a genome of 168,903 bp (34,35) that infects *Escherichia coli*. This phage was selected as it was previously demonstrated to be internalized by mammalian cells, accumulating intracellularly within macropinosomes over time (13,14). Phage lysates were purified and concentrated via ultrafiltration following the Phage on Tap protocol to produce a single, high-titer phage stock that was used for all subsequent assays (32,33). Phage stocks were treated with DNase and RNase to remove extracellular nucleic acids followed by endotoxin removal using 1-Octanol washes. As bacterial endotoxins are known to trigger an innate immune response in TLR4 expressing mammalian cells, we ensured all phage samples were depleted of endotoxin (<1 EU/mL). Despite this, there remained the possibility of bacterial components (i.e., proteins, polysaccharides, nucleic acids) persisting at low levels within our phage lysates. As an additional comparative control, we passed the phage lysate four times through a 0.02 μm filter to generate a phage-free lysate that would also contain any residual bacterial components; henceforth referred to as “Filter control”. As the Filter control contains the same supernatant as the phage lysate but without any phages, it was used as a comparative control to ensure cellular responses were phage-driven and not induced by any bacterial residues or buffer contaminants. We also prepared two additional samples, referred to as “Capsid-only” and “Phage DNA”. The Capsid-only sample was prepared by heating the phages to break the capsid followed by DNase treatment to eliminate the DNA, and thus contained only phage proteins. The Phage DNA sample consisted of the extracted T4 phage genome obtained using the Norgen DNA extraction kit, with DNA integrity checked by T4-specific PCR.

We then investigated whether T4 phages and associated controls could be internalized by our *in vitro* tissue culture model, and whether they activate key intracellular nucleic acid receptors, which stimulate downstream pro-inflammatory immune pathways. T4 phages were applied to both A549 human lung epithelial cells and MDCK-I dog kidney epithelial cell lines and were shown to be rapidly internalized and sequestered within the macropinosomes (Figure 1A & B) (11,12). Once internalized, phages and their nucleic acids could be recognized by the nucleic acid receptor TLR9, a transmembrane protein that resides within endocytic vesicles and preferentially binds DNA from bacteria and viruses. TLR9 is expressed mainly in immune cells (including leukocytes and macrophages) but is also known to be expressed in a range of other cell types, including A549 cells (36). For TRL9 to be activated within the macropinosomes, phage DNA would first have to escape the phage capsid. This could happen through the triggering of the phage ejection apparatus or phage degradation due to acidic conditions found within the endosome (22). Conversely, if the phage particles stay intact, then the phage DNA would not be accessible and TLR9 should not be activated.

**Figure 1:**
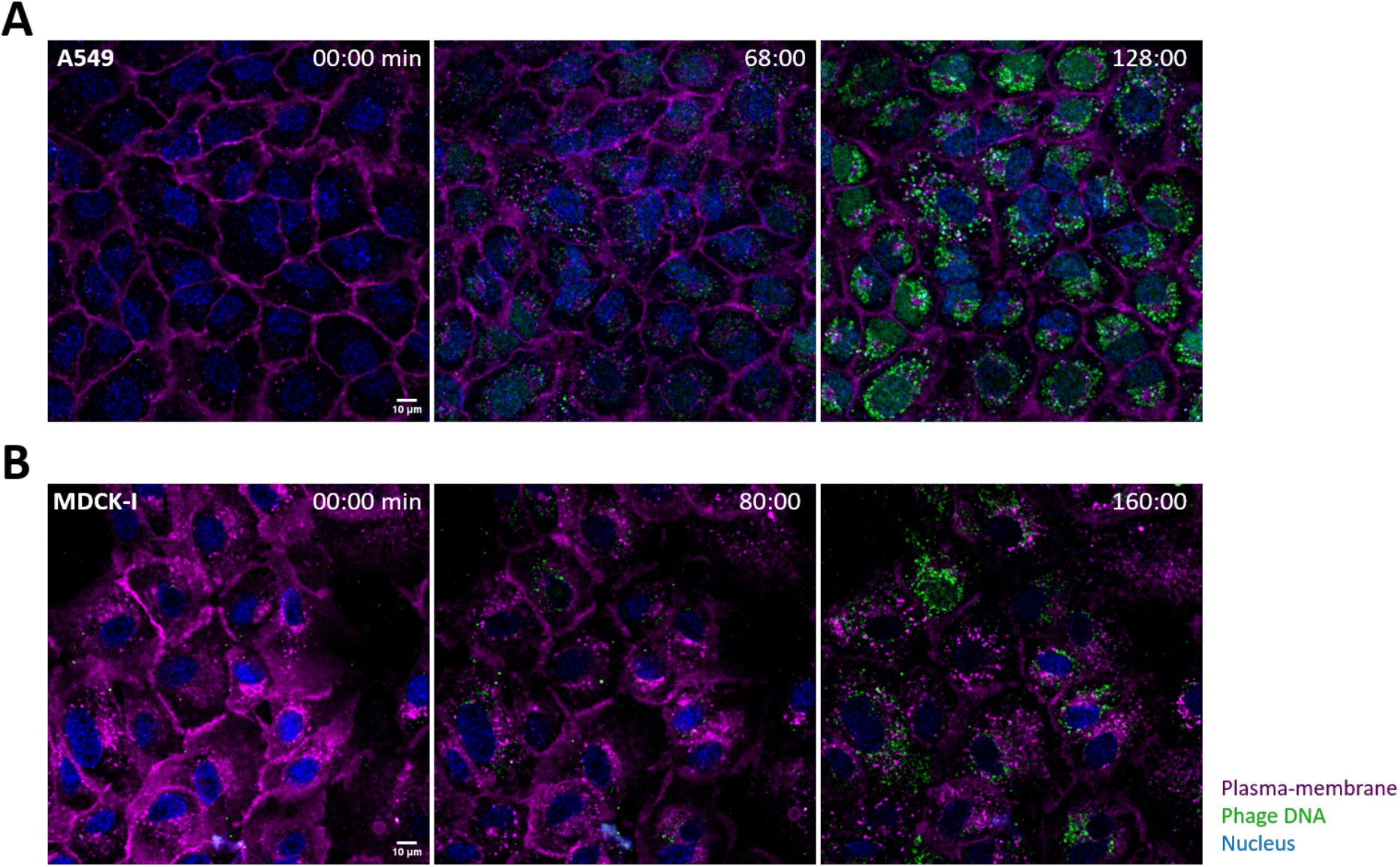
Microscopy images of T4 phage uptake by mammalian cells. **(A)** A549 cells and **(B)** MDCK-I cells incubated with T4 phages for 3h. Images were taken with a confocal microscope every two minutes for three hours. The plasma membrane is shown in magenta, T4 phage DNA in green and the cell nucleus in blue.

Once activated, TLR9 leads to a downstream cascade via the MyD88 pathway resulting in the induction of inflammatory cytokines through activation of NF-*κ*B and other transcription factors, including IRF7, which bind the IFN-β, promoter (37–40). To test this, we used a luciferase-based luminescence assay to detect the downstream activation of TLR9 in activating the IFN-β promoter in A549 cells (Figures 2A & B). We used the reporter plasmid pIFN-β-GL3Luc, which carriers the promoter region of IFN-β gene, and pNF-*κ*B-Luc, which contains five copies of the NF-*κ*B binding motif of the IFN-β promoter, upstream of a luciferase report gene. (41). As positive controls to activate expression from the plasmids we used FLAG-MAVS and pEF-FLAG-RIGI-I(N), which are known to activate the IFN-β promoter, though activation of NF-*κ*B and parallel pathways. One day post-transfection of the reporter plasmid carrying the luciferase cassette, either T4 phage at a titer of 10^9^ PFU/mL or the Filter control, were added to the transfected cells and incubated for two days. Cells were then harvested, lysed and luminescence measured. We saw no activation of luciferase expression from either pNF-kB or pIFN-β-GL30Luc in either the Phage or Filter control samples, while transfection of the positive controls showed strong activation of expression from both plasmids (Figure 2A & B, Supplemental Figure S1). From these results, we conclude that neither NF-*κ*B-dependent activation, nor activation by other elements that activate IFN-ß expression were induced by the internalization of T4 phages. This suggests that the T4 phage capsid remains intact and phage DNA is not exposed nor detected by TLR9 within the macropinosome.

**Figure 2:**
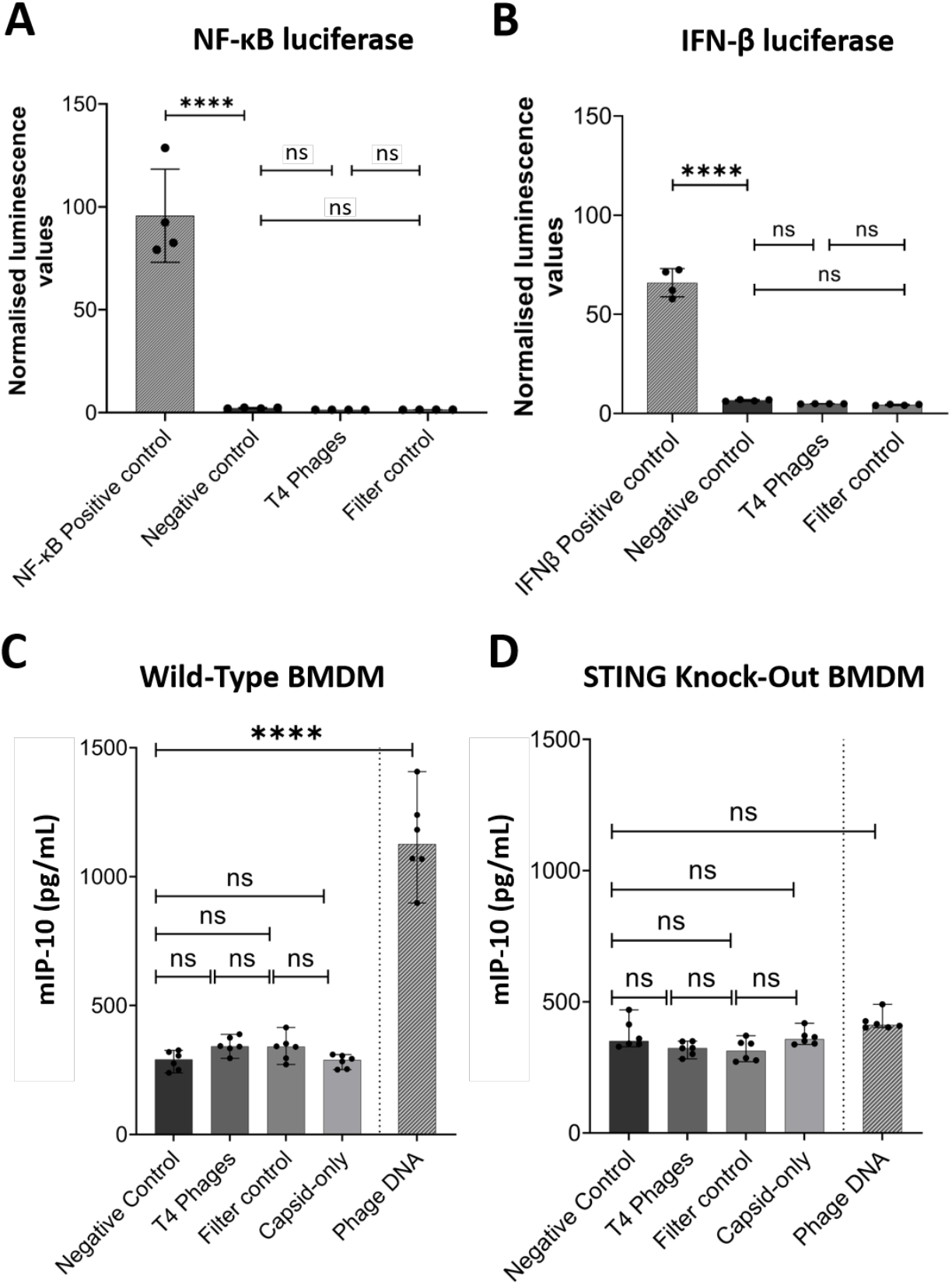
T4 phage do not trigger a pro-inflammatory immune response. A549 cells transfected with either NF-*κ*B-dependent luciferase reporter plasmid **(A)** or IFN-β promoter-dependent luciferase reporter plasmid **(B)** followed by 48 hours incubation with 10^9^ T4 phages/mL or a Filter control. Differentiated WT **(C)** or STING KO **(D)** BMDM cells were incubated for 18 hours 10^7^ T4 phages/mL, Filter control, Capsid-only or transfected with phage DNA using Lipofectamine 2000. P values between the different groups calculated from a one-way ANOVA, shown as stars (P < 0.0001 = ****; A: F (3, 12) = 31.06; B: F (3, 12) = 2.812; C: F (4, 25) = 5.7; D: F (4, 25) = 0.8181).

Following internalization and trafficking, phage particles or DNA may escape the macropinosome and gain access to the cytosol. Here, the presence of free phage DNA would be recognized by the cGAS-STING pathway, leading to the production of IFN-β and inflammatory cytokines (42–45). For phage-mediated activation of this pathway to occur, phage DNA would have to escape both the phage capsid and the macropinosomes to access the cytosol. To test this, we incubated wild-type Bone Marrow-Derived Macrophages (BMDM) or STING knock-out BMDM with either 10^7^ T4 phages/mL, Filter control, or Capsid-only samples for 18 hours. After incubation, the activation of the cGAS-STING pathway was measured by ELISA to measure IFN-β levels (Figures 2C & D). Additionally, as a positive control, we transfected cells with extracted T4 phage DNA using lipofectamine-2000 to demonstrate that phage DNA can activate cGAS-STING. We saw no STING induction in either the Phages, Filter control or Capsid-only samples, while the positive controls transfected with phage DNA showed strong activation of STING in the WT cells (Figure 2C). Comparatively, in the STING-KO BMDM cells, we did not see any activation of the cGAS-STING pathway for any of the controls or samples (Figure 2D). Importantly, both wild-type and STING-KO BMDM cells also have a functional TLR9; suggesting that the fact our STING KO cell lines did not respond to phage stimulation further confirms that TLR9 is indeed not in play. In summary, highly purified T4 phage were internalized by mammalian cells but did not activate pathways downstream of TLR9, including NF-*κ*B-dependent pathways, nor cGAS-STING signaling pathways. This suggests that internalized T4 phages capsids remain intact or are trafficked in such a way as to prevent phage DNA from triggering the innate immune system.

### T4 phage induces protein expression and phosphorylation changes in cell signaling pathways

To investigate the broader cellular responses to phage T4, we utilized an antibody microarray to investigate changes in the expression and phosphorylation of key cell signaling proteins. We used two protein microarrays from Kinexus Biotech, the KAM-1325 microarray that contains 1,325 pan- and phosphosite-specific antibodies covering all the main cellular signaling pathways and can recognise phosphorylated or non-phosphorylated proteins, and the KAM-2000 microarray with 2,000 pan- and phosphosite-specific antibodies. Importantly, the MDCK-I samples were analyzed by the KAM-1325 antibody array, while the A549 samples were analyzed using the improved KAM- 2000 antibody array, which includes most of the antibodies from the prior array, along with 675 additional antibodies for improved detection of cellular changes. We expanded our initial characterization of A549 lung epithelial cells and included a second cell line, the MDCK-I dog kidney epithelial cells, which are known to form strong tight junctions and rapidly internalize and traffic high numbers of T4 phages (Figure 1B) (13,33). Cells were incubated with either purified T4 phages or the Filter control sample for eight hours, followed by cell lysis, chemical cleavage, and protein quantification. MDCK-I cell lysates were directly added to the microarray chip before sending the chip for analysis to Kinexus, whereas the A549 cell lysates were sent to Kinexus where they were applied to the antibody microarrays followed by analysis.

For deconvolution of the microarray datasets, we used the previously developed methodology by Adderley at al. (46). The datasets were mapped onto a network and a pathway analysis, which utilizes random walks to identify chains of phosphorylation events occurring more or less frequently than expected (46). Rather than focusing solely on the largest fold changes, this analysis instead identifies cellular pathway interactions to provide an interpretation of the most important pathways that are influenced by exposure to phages. Briefly, microarray datasets were filtered to remove low signal intensity and/or relatively high error signals compared to control signals. The network analysis was then separately run for both up- and down-regulated phosphorylation events before being assembled into a comparative pathway map (46). Only pathways with more than two intermediates and with fold-changes greater than 5% CFC (% changes from control) were selected for further consideration. This provided us with an overview of the main cellular pathways that were influenced by the presence of phages (Figure 3: MDCK-I and Figure 4: A549). From this analysis, we found 52 hits for the MDCK-I cells, and 150 hits for the A549 cells, which utilized the improved KAM-2000 antibody array. Based on this analysis, we focused our attention on two main pathways – AKT and the CDK1 – that were common across the two antibody microarray datasets.

**Figure 3:**
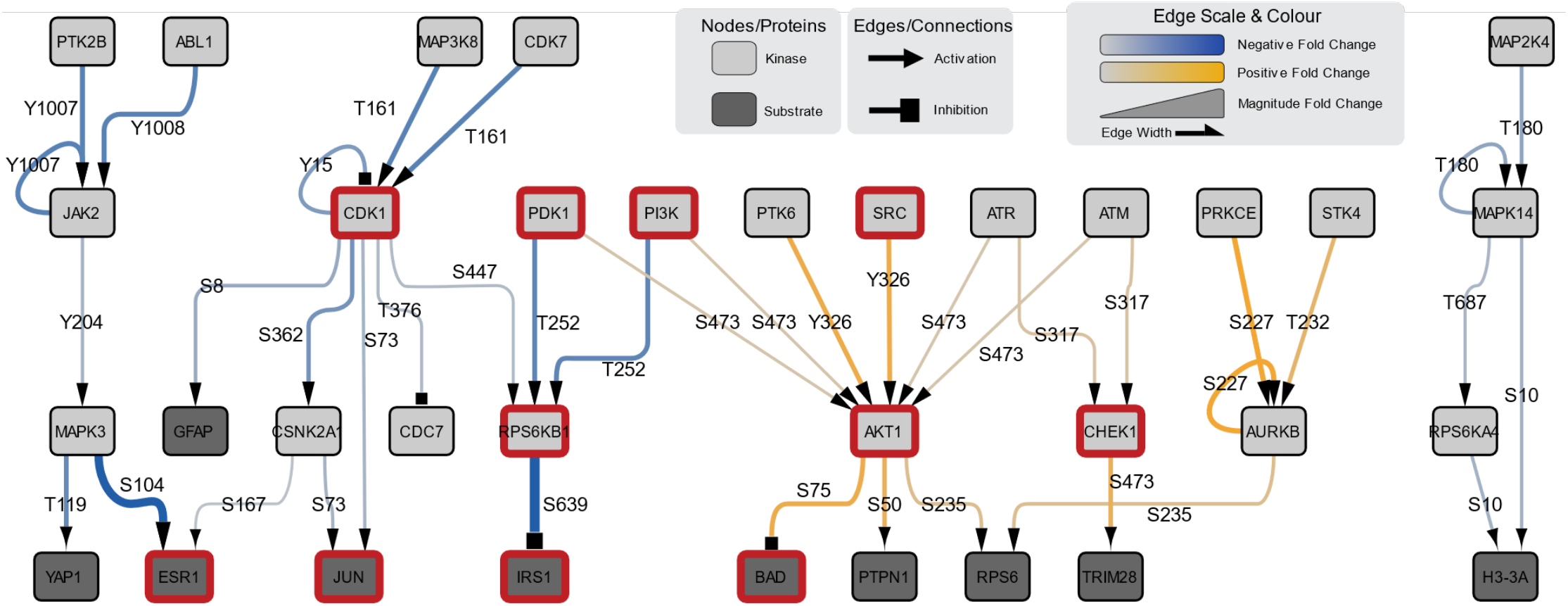
Network protein analysis on MDCK-I cells treated with T4 phages after eight hours of incubation. Kinexus protein microarray with MDCK-I cells after eight hours of incubation with T4 phages. Pathway chart with a detailed pathway of the main leads from the assay. Boxes highlighted in red are proteins discussed in this manuscript.

**Figure 4:**
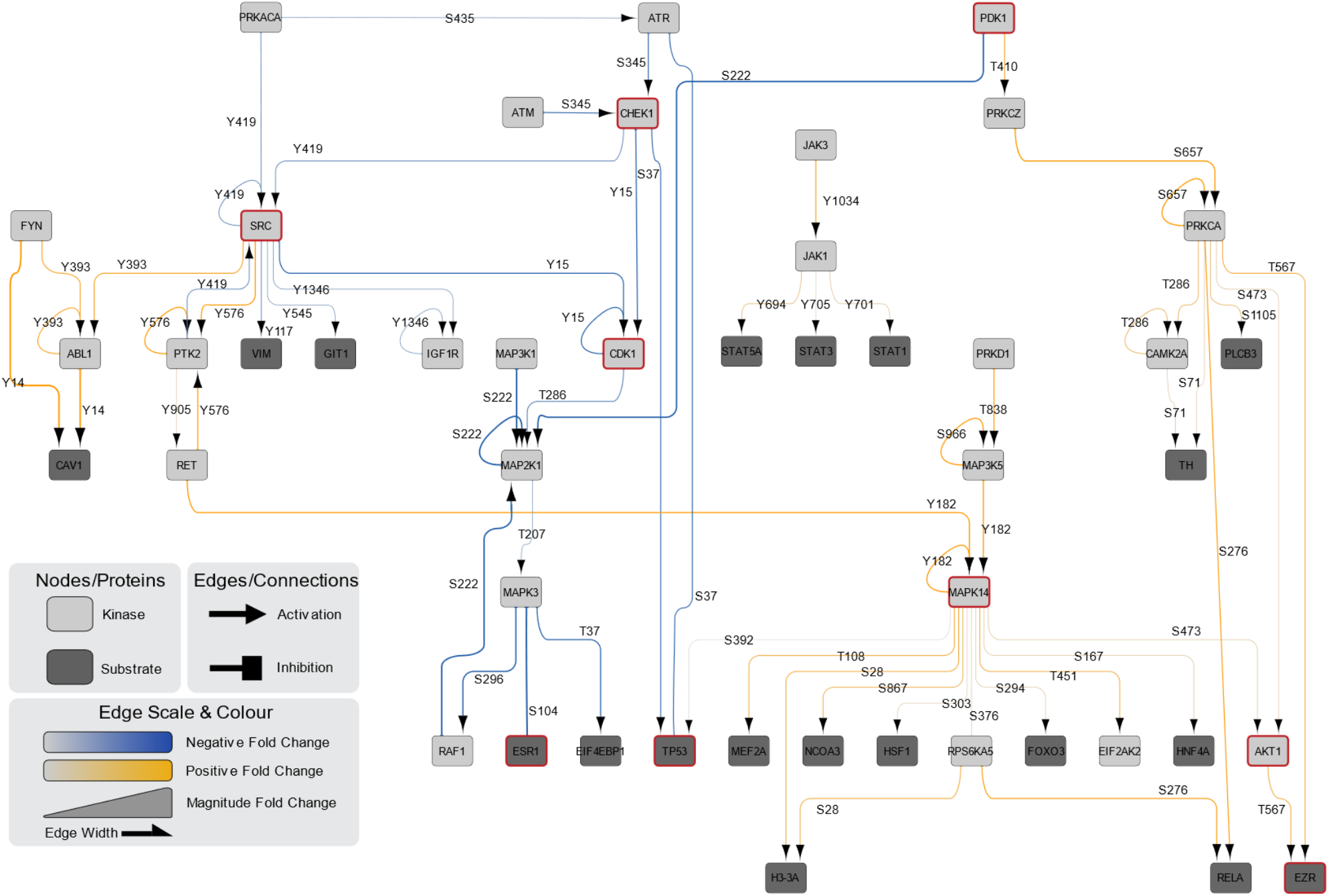
Network protein analysis on A549 cells treated with T4 phages after eight hours of incubation. Kinexus protein microarray with A549 cells after eight hours of incubation with T4 phages. Pathway chart with a detailed pathway of the main up and down-regulated leads from the assay. Boxes highlighted in red are proteins discussed in this manuscript.

### T4 phage activates the AKT pathway promoting cell growth, survival, and macropinocytosis

The AKT signaling pathway is a critical signaling pathway that regulates a myriad of cellular functions, including promoting cell growth, proliferation, survival, and metabolism (47). AKT is a serine/threonine-specific protein kinase that is activated through extracellular growth factors like insulin, which are detected through Receptor Tyrosine Kinases (RTK) or G-Protein Coupled Receptors (GPCR). These receptors recruit PI3K (also called PIK3CA) to the membrane, leading to the recruitment of PDK1 (also called PDPK1), which, once activated, will phosphorylates AKT on T308. Alternatively, PDK1 may recruit mTORC2 which itself will activate AKT through the S473 phosphorylation site (47,48). Once activated, AKT and its downstream effectors will induce a broad range of responses, including glycolysis, protein synthesis, cell survival and proliferation, glycogen synthesis, fatty acid synthesis, and the inhibition of autophagy.

In our MDCK-I datasets, we observed the activation of AKT by PDK1 through the S473 and SRC through the Y326 phosphorylation sites in the presence of T4 phage (Figure 3). Once activated, AKT led to increased inhibition of BAD (BCL2, which is an agonist of cell death) through the S75 phosphorylation site, preventing apoptosis and enhancing cell survival (48–50). Interestingly, in our A549 dataset (Figure 4), we observed similar activation of AKT through the S473 phosphorylation site indirectly by PDK1 but also via MAPK14, which is also known as p38α MAPK. p38α MAPK is activated through environmental stresses and proinflammatory cytokines and usually results in increased cell survival (51). In the A549 cells, the activation of AKT induced the phosphorylation of EZR (Ezrin) through the T567 phosphorylation site (52,53). Importantly, EZR acts as an intermediary between the plasma membrane and the actin cytoskeleton of the cell, with its activation being required for the fusion of cell-to-cell membranes and the formation of endosomes (54). Macropinocytosis, which is the process T4 phages utilize to access the cell, requires significant actin reorganization to generate the membrane ruffles and cell-to-cell membrane fusion to form macropinosomes. As such, the activation of Ezrin through AKT may lead to a positive feedback loop resulting in enhanced phage uptake through the macropinocytosis pathway.

### T4 phage inhibits the CDK1 pathway to delay cell cycle progression and prolong cellular growth

The activity of cyclin-dependent kinases (CDKs) controls all aspects of cell division. CDK1 is implicated in many, if not all, cell cycle regulation pathways and is the central hub for regulating cells progressing through the G2 and mitosis phases of the cell cycle (55–57). CDK1 is essential and sufficient to drive the mammalian cell cycle, including the entry and exit of mitosis and signaling the start of the growth proliferation phase (56,58,59).

In our MDCK-I samples (Figure 3), we observed an in-direct downregulation of CDK1 at the T161 and Y15 phosphorylation sites. The downregulation of CDK1 would inhibit cells from progressing through the G2 and mitotic phases of the cell cycle. This resulted in cascading down-regulation of other cell cycle effectors, including JUN, which is a transcription factor implicated in the prevention of apoptosis and is responsible for the progression of the cell cycle through the G1 phase, via downregulation at the phosphorylation site S73 (60). With the reduction of JUN’s activation, cells’ progression through the G1 phase of the cell cycle would be delayed, keeping cells in a prolonged state of cellular growth. Simultaneously, we observed the downregulation of the activation of Ribosomal protein p70S6K (S6 kinase beta-1 also called RPS6KB1), which can regulate both cell death and proliferation (61). This down-regulation was mediated through two distinct phosphorylation sites, being the T252 phospho-site, which was acted upon by PI3K and PDK1, and S447, which was acted upon by CDK1. Once inhibited, the RPS6KB1 mediated inhibition of IRS-1 (insulin receptor substrate 1) was removed. Previous reports have suggested that IRS-1 can further activate PI3K, thereby (49) leading to a positive feed-back-loop where IRS-1 activates PI3K to further increase AKT activation again (47). Interestingly, we also observed the down-regulated activation of ESR1 (Estrogen Receptor *α*) in both MDCK-I and A549 arrays from the upstream effectors CDK1 and SRC. ESR1 is a large and complex gene that is regulated by multiple elements and encodes for the estrogen receptor and transcription factor, both of which are critical for regulating downstream processes related to cell metabolism, survival, and proliferation (62). Broadly, phages down-regulated key cell cycle effectors responsible for the progression through the G1 phase and modulating regulatory elements associated with cell metabolism and survival.

In the A549 sample (Figure 4), we observed similar down-regulation of CDK1 as in the MDCK-I array, but here through the in-direct phosphorylation of Y15 site by both SRC and CHEK1 (Checkpoint Kinase 1). CHEK1 plays an essential role in cell cycle regulation and DNA damage response (63). CHEK1 further regulates the G1/S transition (along with other cell cycle checkpoints) and is responsible for preventing cells with DNA damage from progressing through the cell cycle (64). At the same time, we saw that CHEK1 was down-regulating the phosphorylation of the tumor suppressor protein TP53 through the S37 phospho-site (65,66). If no damage in the cell DNA were present, TP53 would be rapidly degraded via the proteasome and no build-up of the protein would be observed. If DNA damage were observed in the cell, TP53 would be phosphorylated, and its intracellular concentration would rise, subsequently inducing cell cycle arrest and apoptosis. Thus, phages were down-regulating CHEK1, which led to a subsequent decrease in the activation of CDK1, both of which prevent cells from progressing through the G1 phase of the cell cycle, while also depressing TP53 activation, leading to the prevention of cell cycle arrest and apoptosis.

### Validation of phage-induced cellular changes

From the microarrays, we observed a common pattern where T4 phages induced cell cycle arrest at the G1 phase. We experimentally attempted to see if phage application to A549 cells led to differences in cellular growth and proliferation. Following the application of T4 phage to A549 cells, we quantified the number of cells every 24 hours for four days and a final time point at seven days (Figure 5A). We saw no significant differences in cell proliferation between cells incubated with phages compared to the Filter control. This suggests that the phage-mediated effects on cell cycle arrest and growth were not sufficient to influence aggregate cellular growth and proliferation in this assay. Next, we utilized a comprehensive FACS assay that measured the DNA concentration for each cell to assign them to either a cell cycle phase or cell death. Briefly, A549 cells at 70% confluency were incubated with either T4 phages or the Filter control for eight or 24 hours before collecting cells, treatment with propidium iodide, and analysis via FACS (Figure 5B) (67). While we did not see any significant differences in cell cycle between the phage-treated and Filter control cells at 8-hour time point, we did observe a significant increase in the proportion of phage treated cells in the G0/G1 phase of the cell cycle compared with the Filter control at the 24 hour time point (P = 0.0463). This suggests that, in line with our microarray observations, T4 phage application to A549 cells leads to a prolonged G0/G1 phase that would facilitate broad changes in metabolism, cellular growth, and cell survival.

**Figure 5:**
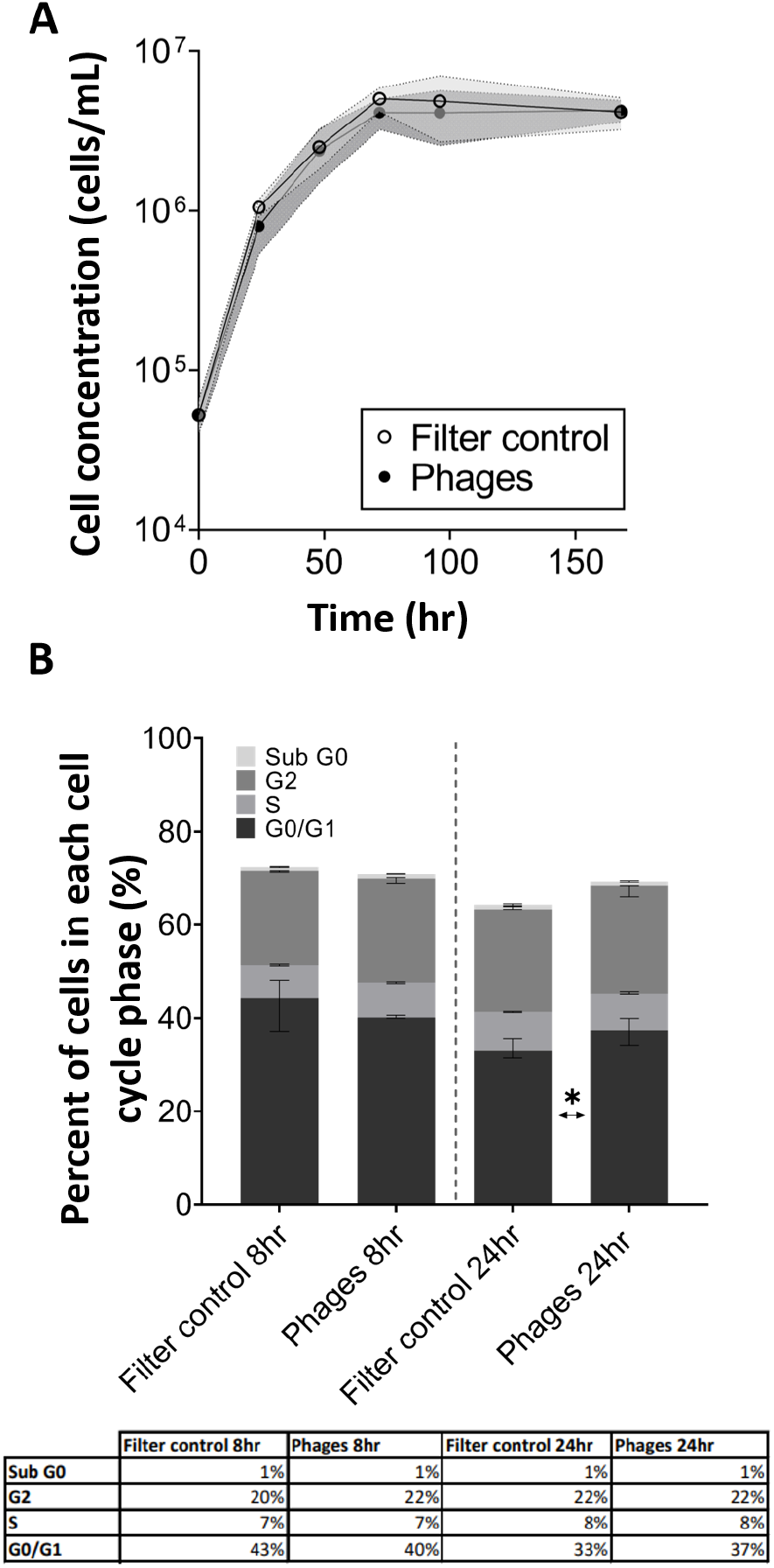
Cell proliferation. **(A)** The proliferation of A549 cells over time with phages in dark circles and the filter control sample in empty circles (shaded area representing 95% CI). **(B)** Cell cycle stage repartition within the A549 cell population after 8 or 24 hr incubation with phages or Filter control (error bars representing 95% CI), including a table presenting the percentage of cells in each cell cycle stage of 3 independent replicates of FACS assay each with 100000 cells analysed. P values of each cell cycle stage between the control and the assay were calculated using a two-way ANOVA, shown as stars (C: F (3, 32) = 2.237).

## DISCUSSION

We observed substantial cellular responses following the application of T4 phages to our *in vitro* cell lines (Figure 6). Importantly, we did not see gene activation from the intracellular DNA-sensing receptors TLR9 and cGAS-STING, suggesting that internalized phages are tightly trafficked to prevent the triggering of the innate immune system. From the antibody microarray results, we observed broad protein phosphorylation responses that revealed common patterns across two cell lines following T4 phage treatment. This suggests that exogenous T4 phages were sensed by cellular receptors (RTK and GPCRs) and internalized by non-specific macropinocytosis. Phages promoted cell survival, proliferation, and metabolism signaling through the activation of the AKT pathway and its downstream effectors. This is consistent with a re-organization of the actin cytoskeleton, which is critical for macropinocytosis and suggestive of a positive feedback loop stimulating further phage uptake. We showed experimentally that phages also prevented cell cycle progression through the downregulation of CDK1 and its effectors. This impaired progression through the G1 phase of the cell cycle and instead left cells in a prolonged state of cellular growth. Overall, these changes suggest that while T4 phages have a benign innate immunological effect on the cell, they do broadly affect cellular response via protein phosphorylation networks. We propose that these *in vitro* mammalian cells are internalizing phages as a food source to maximize their growth and metabolism.

**Figure 6:**
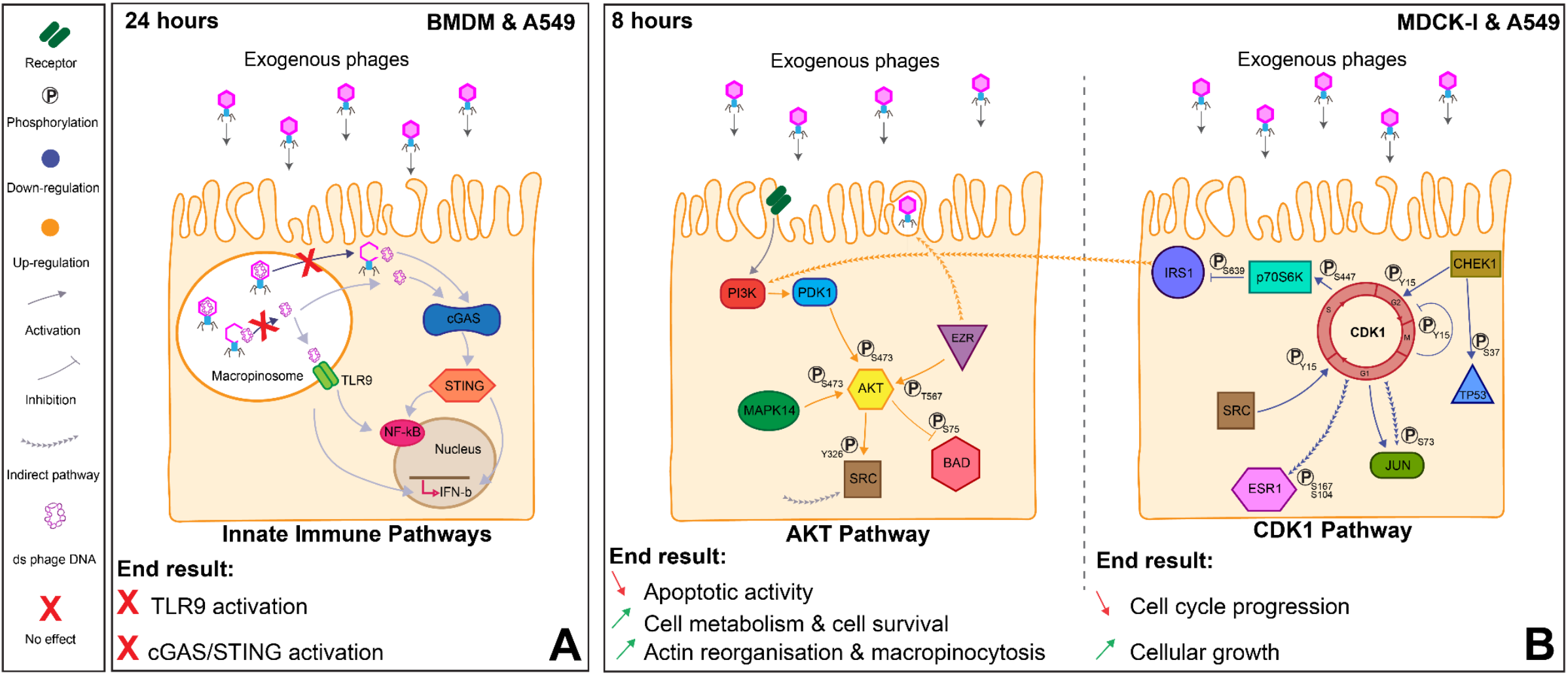
Overview of the effect of exogenous phages on cellular pathways. **(A)** Innate immune pathways in BMDM and A549 cells. Phage DNA is protected by the phage capsid and is not detected by the TLR9 or cGAS-STING. **(B)** The effect of phages on MDCK-I and A549 cells after eight hours. The AKT pathway on the left and the CDK1 pathway on the right show the major cellular changes detected in response to T4 phage.

To activate key inflammatory DNA-sensing innate immune pathways, phage DNA must be accessible in either the cytosol or macropinosomes. We observed that this was rarely the case in mammalian cells treated with T4 phage. Our previous research demonstrated that macropinosome-internalized phages were maintained and accumulated within the cell over time (14). A smaller subset of these internalized macropinosome-bound phages was capable of translocation through the basolateral side of the cell, while others co-localized with lysosomes for further degradation (13). Here we demonstrate that internalized phages did not activate either the TLR9 nor the cGAS-STING pathways, both of which detect dsDNA within the macropinosome, and cytosol respectively, as demonstrated through the lack of downstream activation of NF-*κ*B and IFN-β. This suggests two mechanisms, firstly, that phage uptake and transport by the mammalian cell are tightly regulated by the cell with free phages unlikely to be exposed in the cell cytoplasm, and secondly, phages are not actively degraded or triggered within the macropinosomes, and phage DNA is not exposed to TLR9 receptors (Figure 2). Further experiments are required to explore if different conditions, such as incubation time, pH, temperature, or inflammation state, as well as differences between phages applied or cell lines used, affect the transport and degradation of phages and the subsequent activation of innate immune response and cytokine production (29– 31,68)

We utilized an antibody microarray and a pathway analysis to identify chains of protein phosphorylation events and synergistic interaction networks to decipher cellular pathways of interest (46). From this analysis, we identified two main pathways – AKT and CDK1 – that were affected by T4 phages across our two *in vitro* cell lines. AKT is at the center of a multitude of different cellular processes ranging from the cell cycle, apoptosis, cell survival, glucose metabolism, and the immune system (47,69). The AKT pathway auto-regulates depending on the environmental stress factors, especially in response to the level of extracellular nutrients available for the cell. The activation of AKT at phosphorylation site S473 is known to activate the uptake of glucose for energy production and to promote cellular growth (70), inhibit FoxO proteins to promote anti-apoptotic and cell survival pathways (71,72), and lead to the downstream activation of the Wnt pathway that triggers the entry of cells from G0 into G1 phase (73). Similarly, recent work investigating LNCaP epithelial cells incubated with either T4 phage or M13 filamentous phage found phages mediated the up regulation of the PI3K/AKT pathway leading induced changes in integrin expression as well as increased cell survival (74). We further observed that CDK1 was downregulated by the presence of T4 phages in both cell lines. CDK1 is known to be at the center of all the control checkpoints for the cell cycle and its activation is required for cells to move between cell cycle phases (56). The inhibition of CDK1 led to cascading down-regulation of the cell cycle effector JUN, whose activation is required for the progression of the cell cycle through the G1 phase (60). Simultaneously, we saw a down-regulation of apoptosis via AKT inhibition of BAD and the downregulation of the TP53 phosphorylation (57,75,76). Finally, we further observed AKT activation leading to the up regulation of Ezrin (EZR) at phosphosite T567. The membrane-cytoskeleton linker Ezrin is mainly expressed in epithelial cells and its activation is required for macropinocytosis and for the efficient fusion of vesicles with lysosomes (54,77). Mammalian cells utilized macropinocytosis to take up nutrients in bulk from the extracellular space, with its initiation stimulated by growth factors and PI3K activation, both of which were activated in our cell lines (78). These cellular responses suggest that T4 phage application leads to further uptake and degradation of phages by macropinocytosis, with internalized phages being used as a food source to promote cellular growth.

The most studied group of bacteriophages are the tailed phages (order *Caudoviricetes*) which consist of a complex proteinaceous capsid that houses a highly compressed nucleic acid genome (79). From a macromolecular stance, phages are highly condensed packets of nucleotides in an amino acid shell. Phage T4, which has the most complex capsid structure of any virus, has a capsid mass of 194 MDa (80), of which 55% comprises DNA with a GC content of 34%, while the remaining content consisting of capsid proteins containing all essential amino acids (81). Amino acids are necessary nutrients for the cultivation of mammalian cells *in vitro* with their consumption rate and metabolic flux impacting the growth, metabolism, and regulation of the cell cycle (82). Similarly, nucleotides are required for a wide variety of biological processes, growth, and proliferation, and are constantly synthesized *de novo* in all cells. Nucleotide metabolism supports both RNA synthesis, including ribosomal and messenger RNA to enable biomass production, and DNA replication to enable cell growth and division (83). However, nucleotide production is energy-demanding, and in comparison, with other nutrients, endogenous nucleotide production is favored, as extracellular availability of these nutrients is usually negligible (78). However, the presence of exogenous phage provides the cells with an abundant source of nucleotides accessible via macropinocytosis. This increased uptake of phage-derived nucleotides led to the downregulation of CHEK1 and TP53, both of which are associated with aberrant DNA damage response (64–66). Phage-derived nucleotides led to the effective arrest of cellular growth by accumulating cells at the G1/S phase, which caused excessive cellular growth and the inhibition of cell proliferation that does not affect cell viability (84,85). Our findings support the conclusion that mammalian cells are internalizing T4 phages, which are composed solely of amino acids and nucleotides, as a food source to promote cell growth through a prolonged G1 phase.

We propose a model whereby phages first contact the mammalian cell membrane through diffusive mass transfer resulting in direct phage-cell interactions (13,86). This facilitates the non-specific uptake of cell membrane-associated-phages via macropinocytosis and the accumulation of active phages within membrane-bound vesicles. Internalized phages steadily accumulate and remain functional within the cell for hours to days and are trafficked through diverse pathways (13,14). Phage-containing vesicles are cycled through macropinosomes, fused with lysosomes for degradation, and exocytosed across the basolateral cell membrane. While internalized T4 phages do not trigger inflammatory DNA-sensing immune pathways, they do activate expansive protein phosphorylation cascades. Broadly, these phage-mediated responses may result in increased cell metabolism, cell survival, and further phage uptake via macropinocytosis, while inhibiting autophagy and cell cycle progression through the G1 phase, likely due to the increased supply of phage-derived DNA, leading to increased nucleotide catabolism and a prolonged stage of cellular growth. Further work is required to determine how broad these phage-mediated cellular effects might be. This should include expanding cellular and metabolomics assays on diverse cell types, particularly primary cells, rather than the cancerous cell lines utilized here, which may have a predisposition for enhanced metabolism and growth. Additional phage types and morphologies should be investigated to decipher how cellular uptake and recognition can promote both the non-inflammatory, cellular growth phenotype reported here, versus with the inflammatory phenotype induced by certain phage species (29–31). Open questions remain as to how and why certain phage species trigger aberrant cellular responses (31), whether internalized phages can infect intracellular pathogens (87), how internalized phage particles are degraded and metabolized, and mechanistically how phage-delivered nucleic acids and proteins are accessed by the cell (17,88).

## MATERIAL AND METHODS

### Bacterial stocks and phage stocks

The bacterial strain used in this study was *Escherichia coli B* strain HER 1024, which was cultured in lysogeny broth (LB) media (10 g tryptone, 5 g yeast extract, 10 g NaCl, in 1 liter of distilled water [dH_2_O]) at 37 °C shaking overnight and used to propagate and titer T4 phages supplemented with 10 mM CaCl_2_ and MgSO_4_. T4 phages were cleaned and purified using the Phage on Tap (PoT) protocol (32) and titered up to a concentration of approximately 10^11^ phages/mL to produce a phage stock solution that was used for all experiments. After purification, phages were treated with DNase-I and RNase and then stored in a final solution of SM Buffer (2.0 g MgSO_4_·H_2_O, 5.8 g NaCl, 50 mL of 1M Tris-HCl pH 7.4, dissolved in 1 liter of dH_2_O) at 4 °C.

### Endotoxin removal

The endotoxin removal protocol followed the Phage on Tap (PoT) protocol (32). The phages lysate was cleaned four times with 1-Octanol to remove endotoxins from the lysate, reducing endotoxins from 5734 EU/mL to 167 EU/mL in the final phage stock solution (10^11^ phages/mL) (see also (33). In all experiments, unless otherwise stated, phages were diluted in endotoxin-free buffers to a final concentration of 10^8^ PFU/mL (unless otherwise stated), resulting in an endotoxin concentration below 1 EU/mL.

### Control sample preparation

Using ultra-pure T4 phage lysates we prepared a new comparative control sample. First, in the Filter control sample, the lysate was passed four times through a 0.02 μm filter to remove phage particles from the lysate. The absence of phages was confirmed using a top agar assay at neat dilution with no plaques observed. Second, the capsid-only sample was obtained by breaking the phage capsid using heat treatment. Phages were heated at around 70 °C to break open the capsid and release DNA. The sample was then treated with DNase-I to degrade the phage DNA and only the empty capsids remained. Again, the absence of active phages was tested using plaque assay. Finally, the phage DNA sample was obtained using the Norgen Phage DNA isolation kit (Norgen Cat#46800) following the manufacturer’s instructions and confirmed using T4 phage-specific PCR.

### Cell line stocks

A549 cells were grown in Ham’s F-12K (Kaighn’s) (also called F12-K) (Life Technologies Australia Pty. Ltd) media with 10% Fetal Bovine Serum (FBS) (Life Technologies Australia Pty. Ltd) at 37 °C and 5% CO2 and supplemented with 1% penicillin-streptomycin (Life Technologies Australia Pty. Ltd). MDCK-I cells were grown in Modified Eagle Medium (MEM) (Life Technologies Australia Pty. Ltd) with 10% FBS supplemented with 1% penicillin-streptomycin (Sigma-Aldrich, Australia) at 37 °C and 5% CO_2_.

### Confocal microscopy

For the confocal microscopy experiment, cells were seeded in an IBIDI μ- Slide 8-well glass-bottom slide (DKSH Australia Pty. Ltd). When cells reached 80 to 90% confluency, cells were incubated for 20 min with the respective culture media for each cell line with 5% Hoechst 33342 stain, excitation/emission 115361/497 nm (Life Technologies Australia Pty. Ltd) and 1% CellMask deep red plasma membrane stain, excitation/emission 649/666 nm (Life Technologies Australia Pty. Ltd). After incubation cells were washed three times with Dulbecco’s phosphate-buffered saline (DPBS) and then left in Hank’s Balanced Salt Solution (HBSS) with 1% FBS until acquisition. Purified phages were labelled with 1% SYBR-Gold, excitation/emission 495/537 nm (Life Technologies Australia Pty. Ltd), following the protocol in Bichet et al. 2021b. 200 μL of clean stained phages were in each well containing cells, right before the start of the acquisition (See the detailed protocol in Bichet et al. 2021b). Cells were imaged with HC PL APO 63x/1.40 Oil CS2 oil immersion objective by Leica SP8 confocal microscope on inverted stand with a hybrid detector (HyD) in real-time. HyD detector was used in sequential mode to detect the phages. One image was acquired every two minutes for two hours. Time lapses were created through post-processing using the FIJI software version 2.0.0-rc-68/1.52f (Schindelin et al., 2012). (See the detailed protocol in Bichet et al. 2021b).

### Luciferase assay

A549 cells were plated at 1.5 x 10^5^ cells/mL in 24 well cell culture plates (Corning) for one day. Once cells reached 70% confluency, the cells were co-transfected using Fugene HD transfecting reagent at a 1:3 ratio (Promega) along with pRL-TK *Renilla* (Renilla Luciferase, internal control; Promega) as an internal transfection control. We used the reporter systems pIFN-β-GL3-Luc or pNF-*κ*B-Luc for luciferase (Firefly Luciferase) and the positive control pEF-FLAG-RIGI-I(N) or FLAG-MAVS for IFN-β and NF-*κ*B respectively, or the negative control pUC-18 (empty-vector). A transfection control well was transfected with peGFPC1 instead of pUC-18 to measure the transfection rate with a fluorescent microscope. Each well was transfected in duplicate. One day post-transfection phages at 10^9^ PFU/mL or Filter control were added to the cell layer for two additional days. Cells were then incubated for 30 min at 4 °C slowing rotating with Passive Lysis Buffer (Promega Cat#E1941). After incubation with the lysis buffers, cells were scraped and collected and spun down for 3 min at high speed. The supernatant was collected and kept at - 20 °C until analysis. The values for firefly luciferase activity were normalized to those of *Renilla* luciferase by calculating the ration of firefly to *Renilla* luminescence.

### BMDMs isolation and differentiation

BMDMs were obtained by differentiating isolated bone marrow cells from the femurs of the STING-deficient and matched wild-type control (89). Briefly, bone marrow cells were flushed, washed, and differentiated in a 20% L929 cell-conditioned medium for six days at 37 °C in a 5% CO2, as described previously (90). The use of mouse tissues was approved by the Monash University Animal Ethics Committee (MARP/2018/067).

### cGAS-STING Phage assay

The day before the phage treatment BDMD cells were detached by gently scraping the flasks and plated at 100 000 cells per well in a final volume of 200 μL in a 96 wells plate. The day after 2 μL of 10^7^ phages/mL solution, Filter control or capsid-only samples (or 1.3 ng of phages DNA complexed with lipofectamine 2000 as control) were added to the BMDM cells for another 18 hours. Murine IP-10 production was measured from 100 μL of the supernatant from the BMDMs using Mouse CXCL10/IP-10/CRG-2 Duo Set ELISA (R&D systems, #Dy466) according to the manufacturer’s protocol.

### KAM-1325 Kinexus antibody microarray

Following the protocol described in (33), confluent MDCK-I were incubated with T4 phages or Filter control samples for eight hours at 37 °C and 5% CO_2_. After incubation, the cells were scraped in lysis buffer before sonication. All samples were treated as chemically lysed proteins and followed the recommended protocol by Kinexus. The MDCK-I proteins were quantified using the Bradford protein concentration assay (Thermofisher). The samples were then incubated on the KAM-1325 array before sending the array to Kinexus for analysis.

### KAM-2000 Kinexus antibody microarray

Following the protocol described in (33), confluent A549 cells were incubated with T4 phages or Filter control samples for eight hours at 37 °C and 5% CO_2_. After incubation with the phages or control sample, the cells were scraped in lysis buffer before sonication. All samples were treated as chemically lysed proteins following the recommended protocol by Kinexus. The A549 proteins were quantified using the Bradford protein concentration assay (Thermofisher) and sent to Kinexus to be run on the KAM-2000 array and analyzed.

### Microarray analysis

The analysis of the microarray dataset was performed using the MAPPINGS V1.0 network analysis program developed by (46). There was redundancy between the antibodies tested across the KAM-1325 and −2000 arrays with different antibodies targeting the same protein for more precision (Supplemental Figures S2 & S3 and Supplemental Tables S1 & S2). First, the signals were filtered and all signals below 1000 units were considered as low for the KAM-1325 array and below 500 units for the KAM-2000 array and removed from the assay. Any signal with a high error relative to signal change was disregarded, and any antibody with a higher total signal error across all the arrays compared to the control arrays was disregarded. Next, each unknown substrate effect, where no known biological data was found, was considered as an activation effect for this analysis All nodes (kinases) without a directed edge towards them were the most probable kinase for the downstream phosphorylation event. This was a consequence of the microarray data not reporting which kinase is responsible for each phosphorylation event. Independent positive and negative network analysis were analyzed and in the case of parallel phosphorylation, only the ones with the greater magnitude values were selected and appeared on the pathway map. To ensure the proper termination of each pathway, three options were chosen: 1-If no other path were available after the last kinase or substrate, then the pathway was stopped. 2-If the last phosphorylation had an inhibitory effect then the pathway was stopped as well. 3-Finally, for pathways with only one downstream option available but with no changes from the data sets between the cells incubated with the phages and the Filter control, then a percentage of fold change was assigned for each of these single paths, either between 0-20% for the experimental data-set and 20% for the control ((46)).

### Cell cycle assay

A549 cells were plated in 12 well plates at 5 x 10^4^ cells/mL. Six hours after plating, phages or Filter control samples were added to cells. Using a cell scraper, cells were collected and counted every 24 hours for four days and then one more time seven days after plating. Cells were counted using a Malassez counting slide and a light microscope. Each sample was tested in duplicate and each well was counted twice.

### FACS assay

A549 cells were plated at 8 x 10^4^ cells/mL. The next day, phages at 10^9^ PFU/mL or Filter control samples were added to the cells and incubated for 8 or 24 hours. Cells were then washed twice with PBS; the washes were collected in a 15 mL falcon tube to prevent the loss of dead floating cells and a bias in the analysis. We added trypsin to the wells to collect the cells in the corresponding 15 mL falcon tubes. Cells were quickly centrifuged before adding 1mL of cold PBS. While vortexing, we slowly added 2.3 mL of ice-cold 100% EtOH. Cells were incubated for 40 min at 4 °C. Cells were centrifuged for 5 min at 300 g and resuspended in 500 μL of cold PBS. Cells were again centrifuged for 5 min at 300 g and resuspended in Cell Cycle Buffer (in PBS add 30 μg/mL of PI, 100 μg/mL of RNase A), incubating the cells at RT in the dark for 45 min. The cells were centrifuged for 5 min 300 g and resuspended in 500 μL of cold PBS before being transferred to a 5 mL polystyrene round bottom tube (Corning). Cells were left at 4 °C until FACS analysis following the (67) protocol. The experiment was performed with triplicate wells for each condition and a control well with no PI (Supplemental Figures S4 & S5). Data was generated on a 4 laser Fortessa X-20, manufactured by Becton Dickinson (BD). 100 000 cells were analyzed for each assay and assigned to a cell cycle stage: G0/G1, G2, S or Sub G0.

### Statistics

n represents the number of samples analyzed, each sample was performed in duplicate. All the statistics across this article were done using the GraphPad Prism software using One-way ANOVA or two-way ANOVA. The results of the statistics are represented with stars on top of the corresponding data. All of the microarray experiments were performed with only one sample, no statistics were applicable to these results.

## Acknowledgements & funding

We thank the following labs for kindly providing the cell lines; Hudson Institute of Medical Research and the Oncogenic Signaling Lab for providing the A549 cell line; Stephane Chappaz for preparing the mice femurs and extracting the BMDMs. We thank Rongtuan Lin (McGill University, Canada), Naoto Ito (Gifu University Japan), Natalie Borg (RMIT, Australia) and Takashi Fujita (Kyoto University, Japan) for kindly providing the INF-ß GL3 and NF-*κ*B, pEF-FLAG-RIGI-I(N) and FLAG-MAVS plasmids, respectively. Authors acknowledge and thank the following facilities for kindly providing equipment and guidance: Monash Micro Imaging Facility for help with microscopy acquisition, and the FlowCore for assistance with flow cytometry analysis. Marion C. Bichet was supported by Monash Graduate Scholarship (MGS). This work, including the efforts of Jeremy J. Barr, was funded by the Australian Research Council DECRA Fellowship (DE170100525), National Health and Medical Research Council (NHMRC: 1156588 & 1125704), and the Perpetual Trustees Australia award (2018HIG00007).

## Authors contributions

Conceptualization, MCB, JJB; Methodology, MCB, JA, LA, CDe, CDa, GP, RL, RP, GWM, CD, JJB; Formal Analysis, MCB, JA, LA, LJG, GP; Investigation, MCB; Resources, MPG, GWM, CD, JJB; Writing – Original Draft Preparation, MCB, JJB; Writing – Review and Editing, all authors contributed; Supervision and Funding Acquisition, JJB.

## REFERENCES

1. Barr JJ. A bacteriophages journey through the human body. Immunol Rev. 2017;279(1):106–22.

2. Merril CR. Bacteriophage interactions with higher organisms. Transactions New York Academy of Sciences. 1974;

3. Dabrowska K, Switała-Jelen K, Opolski A, Weber-Dabrowska B, Gorski A. Bacteriophage penetration in vertebrates. J Appl Microbiol [Internet]. 2005 [cited 2020 Apr 15];98(1):7–13.

4. Shkoporov AN, Hill C. Bacteriophages of the Human Gut: The “Known Unknown” of the Microbiome. Vol. 25, Cell Host and Microbe. Elsevier; 2019. p. 195–209.

5. Clokie MR, Millard AD, Letarov A V, Heaphy S. Phages in nature. Bacteriophage. 2011 Jan;1(1):31–45.

6. Sender R, Fuchs S, Milo R, Berg RD, Bianconi E, Piovesan A, et al. Are We Really Vastly Outnumbered? Revisiting the Ratio of Bacterial to Host Cells in Humans. Cell. 2016 Jan;164(3):337–40.

7. Stilling RM, Dinan TG, Cryan JF. Microbial genes, brain & behaviour-epigenetic regulation of the gut-brain axis. Genes Brain Behav. 2014;13:69–86.

8. Shkoporov AN, Clooney AG, Sutton TDS, Ryan FJ, Daly KM, Nolan JA, et al. The Human Gut Virome Is Highly Diverse, Stable, and Individual Specific. Cell Host Microbe. 2019 Oct 9;26(4):527–541.e5.

9. Shkoporov AN, Stockdale SR, Lavelle A, Kondova I, Heuston C, Upadrasta A, et al. Viral biogeography of the mammalian gut and parenchymal organs. Nat Microbiol. 2022 Aug 1;7(8):1301–11.

10. Liang G, Zhao C, Zhang H, Mattei L, Sherrill-Mix S, Bittinger K, et al. The stepwise assembly of the neonatal virome is modulated by breastfeeding. Nature. 2020 May 28;581(7809):470–4.

11. Huh H, Wong S, St. Jean J, Slavcev R. Bacteriophage interactions with mammalian tissue: Therapeutic applications. Adv Drug Deliv Rev. 2019;145:4–17.

12. Dabrowska K, Switała-Jelen K, Opolski A, Weber-Dabrowska B, Gorski A. Bacteriophage penetration in vertebrates. J Appl Microbiol. 2005 Jan;98(1):7–13.

13. Nguyen S, Baker K, Padman BS, Patwa R, Dunstan RA, Weston TA, et al. Bacteriophage transcytosis provides a mechanism to cross epithelial cell layers. Racaniello VR, editor. mBio [Internet]. 2017 Nov 21;8(6):1–14.

14. Bichet MC, Chin WH, Richards W, Lin YW, Avellaneda-Franco L, Hernandez CA, et al. Bacteriophage uptake by mammalian cell layers represents a potential sink that may impact phage therapy. iScience. 2021;24(4):102287.

15. Górski A, Wazna E, Dabrowska BW, Dabrowska K, Switała-Jeleń K, Miedzybrodzki R. Bacteriophage translocation. FEMS Immunol Med Microbiol. 2006 Apr 1;46(3):313–9.

16. Lehti TA, Pajunen MI, Skog MS, Finne J. Internalization of a polysialic acid-binding Escherichia coli bacteriophage into eukaryotic neuroblastoma cells. Nat Commun. 2017 Dec 4;8(1):1915.

17. Tao P, Mahalingam M, Marasa BS, Zhang Z, Chopra AK, Rao VB. In vitro and in vivo delivery of genes and proteins using the bacteriophage T4 DNA packaging machine. Proc Natl Acad Sci U S A. 2013 Apr 9;110(15):5846–51.

18. King JS, Kay RR. The origins and evolution of macropinocytosis. Vol. 374, Philosophical Transactions of the Royal Society B: Biological Sciences. Royal Society Publishing; 2019.

19. Lim JP, Gleeson PA. Macropinocytosis: an endocytic pathway for internalising large gulps. Immunol Cell Biol. 2011;89:836–43.

20. Kerr MC, Teasdale RD. Defining Macropinocytosis. Traffic. 2009 Apr;10(4):364–71.

21. Falcone S, Cocucci E, Podini P, Kirchhausen T, Clementi E, Meldolesi J. Macropinocytosis: Regulated coordination of endocytic and exocytic membrane traffic events. J Cell Sci. 2006 Nov;119(22):4758–69.

22. Buckley CM, King JS. Drinking problems: mechanisms of macropinosome formation and maturation. FEBS J. 2017;284:3778–90.

23. Schlee M, Hartmann G. Discriminating self from non-self in nucleic acid sensing. Nature. 2016 [cited 2021 Oct 24];16.

24. Van Belleghem JD, Dabrowska K, Vaneechoutte M, Barr JJ. Phage interaction with the mammalian immune system. In: Phage Therapy: A Practical Approach. Springer International Publishing; 2019. p. 91–122.

25. Górski A, Kniotek M, Perkowska-Ptasińska A, Mróz A, Przerwa A, Gorczyca W, et al. Bacteriophages and Transplantation Tolerance. Transplant Proc. 2006 Jan;38(1):331–3.

26. Gorski A, Dabrowska K, Switala-Jeleń K, Nowaczyk M, Weber-Dabrowska B, Boratynski J, et al. New insights into the possible role of bacteriophages in host defense and disease. Med Immunol. 2003 Feb;2(1):2.

27. Focà A, Liberto MC, Quirino A, Marascio N, Zicca E, Pavia G. Gut inflammation and immunity: What is the role of the human gut virome? Mediators Inflamm. 2015;2015.

28. Duerkop BA, Hooper L V. Resident viruses and their interactions with the immune system. Nat Immunol. 2013 Jul;14(7):654–9.

29. Gogokhia L, Buhrke K, Bell R, Hoffman B, Brown DG, Hanke-Gogokhia C, et al. Expansion of Bacteriophages Is Linked to Aggravated Intestinal Inflammation and Colitis. Cell Host Microbe. 2019 Feb 13 [cited 2020 Sep 29];25(2):285–299.e8.

30. Sweere JM, van Belleghem JD, Ishak H, Bach MS, Popescu M, Sunkari V, et al. Bacteriophage trigger antiviral immunity and prevent clearance of bacterial infection. Science (1979). 2019 Mar 29 cited 2020 Apr 20];363(6434).

31. Adiliaghdam F, Amatullah H, Digumarthi S, Saunders TL, Rahman RU, Wong LP, et al. Human enteric viruses autonomously shape inflammatory bowel disease phenotype through divergent innate immunomodulation. Sci Immunol. 2022 Apr 1;7(70).

32. Bonilla N, Rojas MIMI, Cruz GNF, Hung SHSH, Rohwer F, Barr JJ. Phage on tap–a quick and efficient protocol for the preparation of bacteriophage laboratory stocks. PeerJ. 2016 Jul 26;2016(7):e2261.

33. Bichet MC, Patwa R, Barr JJ. Protocols for studying bacteriophage interactions with in vitro epithelial cell layers. STAR Protoc. 2021 Sep 17;2(3):100697.

34. Miller ES, Kutter E, Mosig G, Arisaka F, Kunisawa T, Rüger W. Bacteriophage T4 genome. Microbiology and molecular biology reviews. 2003 Mar;67(1):86–156, table of contents.

35. Subedi D, Barr JJ. Temporal Stability and Genetic Diversity of 48-Year-Old T-Series Phages. mSystems. 2021 Feb 23;6(1).

36. Droemann D, Albrecht D, Gerdes J, Ulmer AJ, Branscheid D, Vollmer E, et al. Human lung cancer cells express functionally active Toll-like receptor 9. Respir Res. 2005 Jan;6(1):1–10.

37. Wagner H. The immunobiology of the TLR9 subfamily. Trends Immunol. 2004;25(7).

38. Marongiu L, Gornati L, Artuso I, Zanoni I, Granucci F. Below the surface: The inner lives of TLR4 and TLR9. J Leukoc Biol. 2019;(106):140–60.

39. Huang X, Yang Y. Targeting the TLR9MyD88 pathway in the regulation of adaptive immune responses. Vol. 14, Expert Opinion on Therapeutic Targets. Taylor & Francis; 2010. p. 787–96.

40. Kumagai Y, Takeuchi O, Akira S. TLR9 as a key receptor for the recognition of DNA. Vol. 60, Advanced Drug Delivery Reviews. Elsevier; 2008. p. 795–804.

41. Lin R, Génin P, Mamane Y, Hiscott J. Selective DNA Binding and Association with the CREB Binding Protein Coactivator Contribute to Differential Activation of Alpha/Beta Interferon Genes by Interferon Regulatory Factors 3 and 7. Mol Cell Biol. 2000 Sep;20(17):6342–53.

42. Chen Q, Sun L, Chen ZJ. Regulation and function of the cGAS-STING pathway of cytosolic DNA sensing. Nat Immunol. 2016;17(10):1142–9.

43. Tan X, Sun L, Chen J, Chen ZJ. Detection of Microbial Infections Through Innate Immune Sensing of Nucleic Acids. Annu Rev Microbiol. 2018 Sep 8;72(1):447–78.

44. Motwani M, Pesiridis S, Fitzgerald KA. DNA sensing by the cGAS–STING pathway in health and disease. Nat Rev Genet. 2019;20.

45. Kwon J, Bakhoum SF, Kettering S. The Cytosolic DNA-Sensing cGAS-STING Pathway in Cancer. aacrjournals.org Cancer Discov. 2020;10:26–39.

46. Adderley J, O’donoghue F, Davis S, Doerig C. MAPPINGS v1.0, a tool for network analysis of large phospho-signalling datasets: application to host erythrocyte response to Plasmodium infection. Res Sq. 2021 Sep 13 [cited 2021 Oct 26];

47. Manning BD, Cantley LC. AKT/PKB Signaling: Navigating Downstream. Cell. 2007 Jun 29;129(7):1261–74.

48. Cantley LC. The Phosphoinositide 3-Kinase Pathway. Science (1979). 2002 May 31 [cited 2021 Aug 27];296(5573):1655–7.

49. Del Peso L, González-García M, Page C, Herrera R, Nuñez G. Interleukin-3-Induced Phosphorylation of BAD Through the Protein Kinase Akt. Genome Issue. 1997;278(5338):687–9.

50. Datta SR, Dudek H, Tao X, Masters S, Fu H, Gotoh Y, et al. Akt Phosphorylation of BAD Couples Survival Signals to the Cell-Intrinsic Death Machinery. Cell. 1997;91:231–41.

51. Cuadrado A, Nebreda AR. Mechanisms and functions of p38 MAPK signalling. Vol. 429, Biochemical Journal. Portland Press; 2010. p. 403–17.

52. Quan C, Sun J, Lin Z, Jin T, Dong B, Meng Z, et al. Ezrin promotes pancreatic cancer cell proliferation and invasion through activating the Akt/mTOR pathway and inducing YAP translocation. Cancer Manag Res. 2019;11:6553–66.

53. Song Y, Ma X, Zhang M, Wang M, Wang G, Ye Y, et al. Ezrin Mediates Invasion and Metastasis in Tumorigenesis: A Review. Front Cell Dev Biol. 2020;8(November):1–12.

54. Marion S, Hoffmann E, Holzer D, le Clainche C, Martin M, Sachse M, et al. Ezrin Promotes Actin Assembly at the Phagosome Membrane and Regulates Phago-Lysosomal Fusion. Traffic. 2011 Apr;12(4):421–37.

55. Squire CJ, Dickson JM, Ivanovic I, Baker EN. Structure and Inhibition of the Human Cell Cycle Checkpoint Kinase, Wee1A Kinase: An AtypicalTyrosine Kinase with a Key Role in CDK1 Regulation. Structure. 2005 Apr;13(4):541–50.

56. Santamaría D, Barrière C, Cerqueira A, Hunt S, Tardy C, Newton K, et al. Cdk1 is sufficient to drive the mammalian cell cycle. Nature. 2007;448.

57. Potapova TA, Daum JR, Byrd KS, Gorbsky GJ. Fine Tuning the Cell Cycle: Activation of the Cdk1 Inhibitory Phosphorylation Pathway during Mitotic Exit. Mol Biol Cell. 2009;20:1737–48.

58. Doree M. Control of M-phase by maturation-promoting factor. Curr Opin Cell Biol. 1990;2:269–73.

59. Hunt T. Maturation promoting factor, cyclin and the control of M-phase. Curr Opin Cell Biol. 1989;1:268–74.

60. Pawlonka J, Rak B, Ambroziak U. The regulation of cyclin D promoters – review. Cancer Treat Res Commun. 2021 Jan;27:100338.

61. Sridharan S, Basu A. Molecular Sciences Distinct Roles of mTOR Targets S6K1 and S6K2 in Breast Cancer. Int J Mol Sci. 2020;21(1199).

62. Lung DK, Reese RM, Alarid ET. Intrinsic and Extrinsic Factors Governing the Transcriptional Regulation of ESR1. Vol. 11, Hormones and Cancer. Springer; 2020. p. 129–47.

63. Patil M, Pabla N, Dong Z. Checkpoint kinase 1 in DNA damage response and cell cycle regulation. Vol. 70, Cellular and Molecular Life Sciences. 2013. p. 4009–21.

64. Sanchez Y, Wong C, Thoma RS, Richman R, Wu Z, Piwnica-Worms H, et al. Conservation of the Chk1 Checkpoint Pathway in Mammals: Linkage of DNA Damage to Cdk Regulation Through Cdc25. Science (1979). 1997;277(5331):1497–501.

65. Brooks CL, Gu W. Ubiquitination, phosphorylation and acetylation: the molecular basis for p53 regulation. Curr Opin Cell Biol. 2003;15:164–71.

66. Kruse JP, Gu W. Modes of p53 Regulation. Cell. 2009;137(15).

67. Crowley LC, Chojnowski G, Waterhouse NJ. Measuring the DNA Content of Cells in Apoptosis and at Different Cell-Cycle Stages by Propidium Iodide Staining and Flow Cytometry. Cold Spring Harb Protoc. 2016 Oct;

68. van Belleghem JD, Clement F, Merabishvili M, Lavigne R, Vaneechoutte M. Pro-and anti-inflammatory responses of peripheral blood mononuclear cells induced by Staphylococcus aureus and Pseudomonas aeruginosa phages. Sci Rep. 2017 Dec 1;7(1).

69. Yang WL, Wu CY, Wu J, Lin HK. Regulation of Akt signaling activation by ubiquitination. Cell Cycle. 2010;9(3).

70. Kumar A, Lawrence JC, Jung DY, Ko HJ, Keller SR, Kim JK, et al. Fat cell-specific ablation of rictor in mice impairs insulin-regulated fat cell and whole-body glucose and lipid metabolism. Diabetes. 2010 Jun;59(6):1397–406.

71. Young ARJ, Narita M, Ferreira M, Kirschner K, Sadaie M, Darot JFJ, et al. Autophagy mediates the mitotic senescence transition. Genes Dev. 2009 Apr 1;23(7):798–803.

72. Guan H, Song L, Cai J, Huang Y, Wu J, Yuan J, et al. Sphingosine kinase 1 regulates the Akt/FOXO3a/Bim pathway and contributes to apoptosis resistance in glioma cells. PLoS One. 2011;6(5).

73. Vadlakonda L, Pasupuleti M, Pallu R. Role of PI3K-AKT-mTOR and Wnt signaling pathways in transition of G1-S phase of cell cycle in cancer cells. Front Oncol. 2013;3 APR.

74. Sanmukh SG, Santos NJ, Barquilha CN, Cucielo MS, Carvalho M de, Reis PP dos, et al. Bacteriophages M13 and T4 Increase the Expression of Anchorage-Dependent Survival Pathway Genes and Down Regulate Androgen Receptor Expression in LNCaP Prostate Cell Line. Viruses. 2021;13(1754).

75. Ayeni JO, Campbell SD. “Ready, Set, Go”: Checkpoint regulation by Cdk1 inhibitory phosphorylation. Fly (Austin). 2014;

76. Santamaría D, Barrière C, Cerqueira A, Hunt S, Tardy C, Newton K, et al. Cdk1 is sufficient to drive the mammalian cell cycle. Nature. 2007;448.

77. Angelo RD’, Aresta S, Blangy A, Del L, Louvard D, Arpin M. Interaction of Ezrin with the Novel Guanine Nucleotide Exchange Factor PLEKHG6 Promotes RhoG-dependent Apical Cytoskeleton Rearrangements in Epithelial Cells. Mol Biol Cell. 2007;18:4780–93.

78. Zhu J, Thompson CB. Metabolic regulation of cell growth and proliferation. Vol. 20, Nature Reviews Molecular Cell Biology. Nature Publishing Group; 2019. p. 436–50.

79. Yap ML, Rossmann MG. Structure and function of bacteriophage T4. Vol. 9, Future Microbiology. Future Medicine Ltd.; 2014. p. 1319–37.

80. Fokine A, Chipman PR, Leiman PG, Mesyanzhinov V v, Rao VB, Rossmann MG. Molecular architecture of the prolate head of bacteriophage T4 [Internet]. 2004. Available from: www.pnas.orgcgidoi10.1073pnas.0400444101

81. Bancbofrt FC, Freifelder D. Molecular Weights of Coliphages and Coliphage DNA I. Measurement of the Molecular Weight of Bacteriophage T7 by High-speed Equilibrium Centrifugation. Vol. 54, J. Mol. Biol. 1970.

82. Salazar A, Keusgen M, von Hagen J. Amino acids in the cultivation of mammalian cells. Vol. 48, Amino Acids. Springer-Verlag Wien; 2016. p. 1161–71.

83. Lane AN, Fan TWM. Regulation of mammalian nucleotide metabolism and biosynthesis. Vol. 43, Nucleic Acids Research. Oxford University Press; 2015. p. 2466–85.

84. Carvalhal A v., Sá Santos S, Calado J, Haury M, Carrondo MJT. Cell growth arrest by nucleotides, nucleosides and bases as a tool for improved production of recombinant proteins. In: Biotechnology Progress. 2008. p. 69–83.

85. Diehl FF, Miettinen TP, Elbashir R, Nabel CS, Darnell AM, Do BT, et al. Nucleotide imbalance decouples cell growth from cell proliferation. Nat Cell Biol. 2022 Aug 1;24(8):1252–64.

86. Barr JJ, Auro R, Furlan M, Whiteson KL, Erb ML, Pogliano J, et al. Bacteriophage adhering to mucus provide a non-host-derived immunity. Proceedings of the National Academy of Sciences. 2013;110(26):10771–6.

87. Møller-Olsen C, Ross T, Leppard KN, Foisor V, Smith C, Grammatopoulos DK, et al. Bacteriophage K1F targets Escherichia coli K1 in cerebral endothelial cells and influences the barrier function. Sci Rep. 2020 Dec 1;10(1).

88. Geier MR, Trigg ME, Merril CR. Fate of bacteriophage lambda in Non-immune germ-free mice. Nature. 1973;246(5430):221–3.

89. Jin L, Hill KK, Filak H, Mogan J, Knowles H, Zhang B, et al. MPYS Is Required for IFN Response Factor 3 Activation and Type I IFN Production in the Response of Cultured Phagocytes to Bacterial Second Messengers Cyclic-di-AMP and Cyclic-di-GMP. The Journal of Immunology. 2011;187(5):2595–601.

90. Ferrand J, Gantier MP. Assessing the Inhibitory Activity of Oligonucleotides on TLR7 Sensing. Methods in Molecular Biology. 2016 Jan;1390:79–90.

