## Supplemental Information for "Mammalian cells internalize bacteriophages and utilize them as a food source to enhance cellular growth and survival"

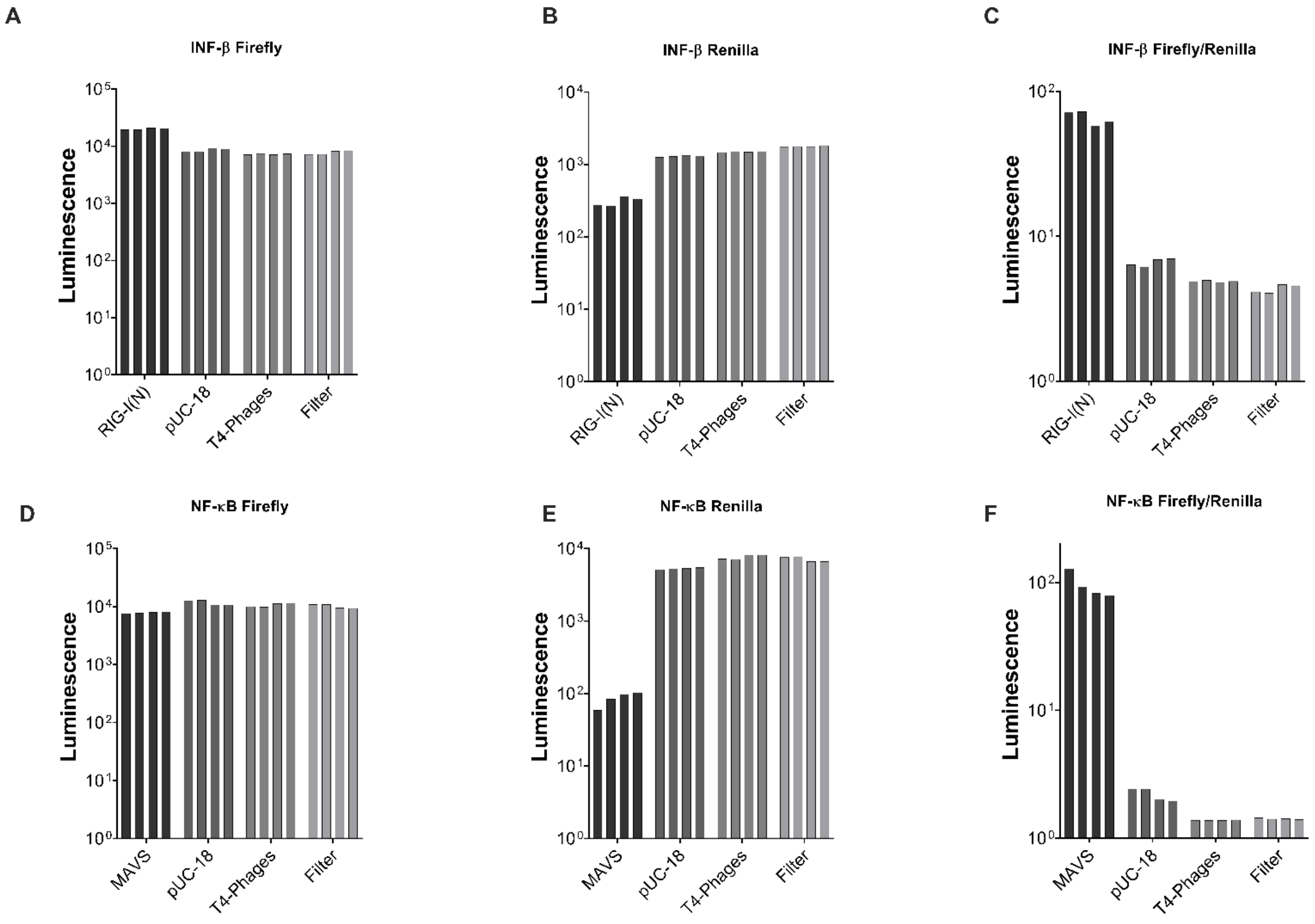


**Figure S1: Luciferase analysis on A549 cells and T4 phages.** Individual results from the luciferase assay with IFN-ß and NF-$\kappa$B from the Firefly and Renilla reporter. **(A)** Firefly IFN-ß luminescence values from each well. **(B)** Renilla IFN-ß luminescence values from each well. **(C)** IFN-ß normalised luminescence values, Firefly/Renilla, from each well. **(D)** Firefly NF-$\kappa$B luminescence values from each well. **(E)** Renilla NF-$\kappa$B luminescence values from each well. **(F)** NF-$\kappa$B normalised luminescence values, Firefly/Renilla, from each well.

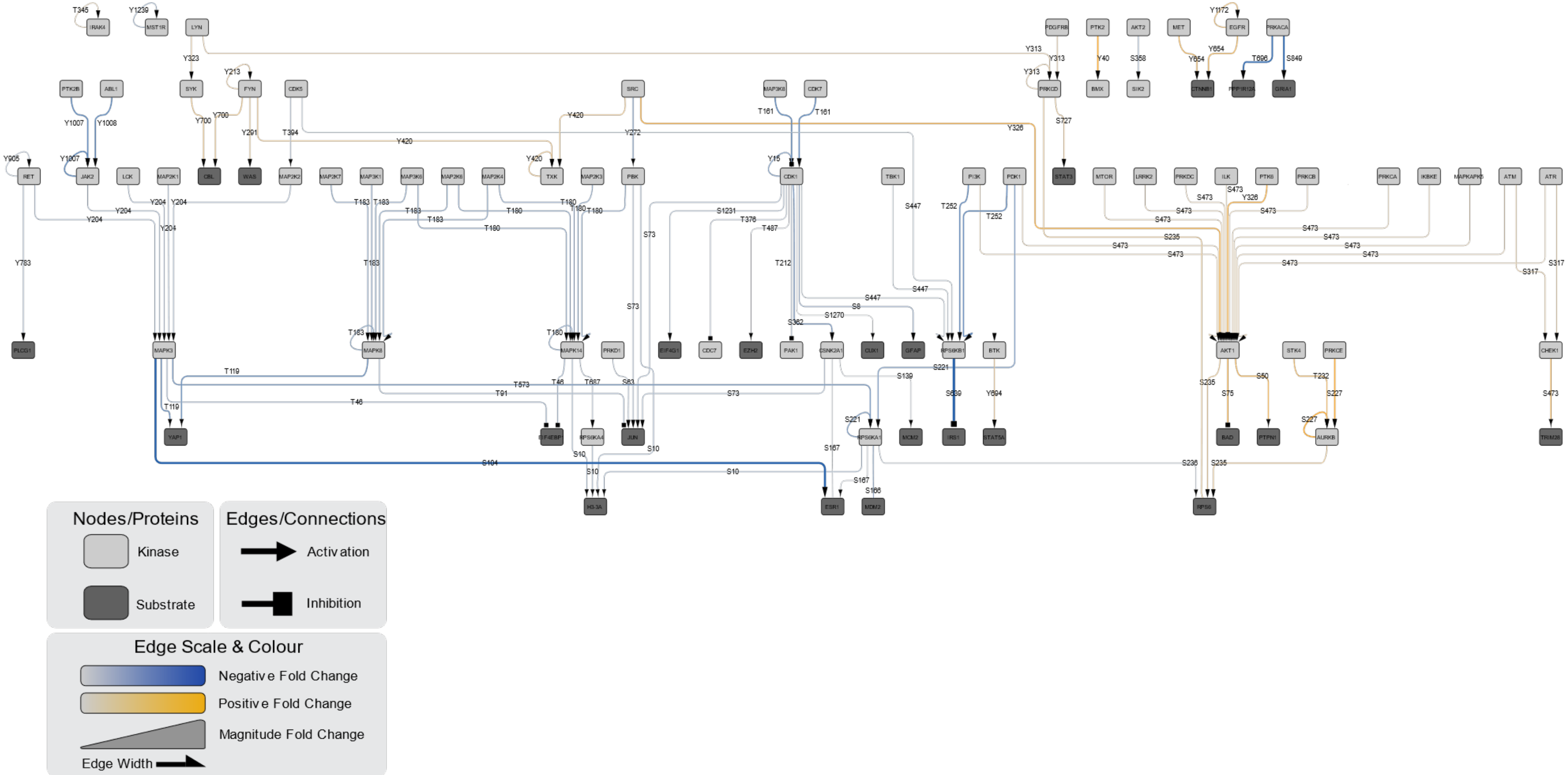


**Figure S2: Microarray map for the MDCK-I sample.** Showing all the interactions detected in the Microarray via the analysis

**Table S1: Table listing the mains leads for the microarray MDCK-I sample.**

| Antibody ID No. | Target Name | Antibody P-Site | % CFC |
|---|---|---|---|
| NN430-1 | Myc | Pan-specific | 371 |
| PN274 | STAT4 | S721 | 179 |
| PN538 | STAM2 | Y374 | 139 |
| PK833 | TRIM28 (TIF1B) | S473 | 116 |
| PK531 | AurKB (Aurora B, AIM-1) | T232 | 93 |
| NK284-1 | PBK | Pan-specific | 87 |
| PN671 | STAT5A | Y694 | 81 |
| PK607 | EphA2 | Y772 | 80 |
| NK273-1 | IRR (INSRR) | Pan-specific | 66 |
| PK817 | SMG1 | T3550 | 54 |
| NK120-7 | p38a MAPK (MAPK14) | Pan-specific | 53 |
| NK121-3 | p38d MAPK (MAPK13) | Pan-specific | 45 |
| PN501 | ACTB | Y53 | 44 |
| NK121-2 | p38d MAPK (MAPK13) | Pan-specific | 40 |
| PK670 | JNK1 (MAPK8) | Y185 | 39 |
| PN655 | SIN3A | S832 | 36 |
| NK181-3 | TYK2 | Pan-specific | 32 |
| PK536 | GRK2 (BARK1, ADRBK1) | S670 | 32 |
| PK836 | TRIM33 (TIF1G) | S1119 | 31 |
| NK250-2 | SIK3 (QSK) | Pan-specific | 28 |
| PK665 | IRAK4 | T345+S346 | 26 |
| NK076-5 | IKKb (IkBKB) | Pan-specific | 23 |
| PN638 | p53 (TP53) | T18+S20 | 20 |
| PK745 | p70S6K (S6Ka, RPS6KB1) | T412 | 14 |
| PK786 | PRP4K (PRP4, PRPF4B) | Y849 | 11 |
| PK645 | Fyn | Y531 | -5 |
| NK255-3 | WNK4 (PRKWNK4) | Pan-specific | -8 |
| PP505 | PPP2CB | T304 | -10 |
| NK156-6 | Raf-B (BRaf) | Pan-specific | -10 |
| NK269-2 | Frk | Pan-specific | -11 |
| PN644 | PPARg-1 | S112 | -12 |
| PN583 | ERa (ESR1) | S167 | -13 |
| NK107-4 | MEKK1 (MAP3K1) | Pan-specific | -13 |
| NK026-6 | CDK2 | Pan-specific | -15 |
| PN745 | CTNNB1 | Y489 | -15 |
| PK879 | ERK1 (MAPK3) | S283 | -15 |
| PK608 | EphA3 | Y779 | -16 |

| | | | |
|---|---|---|---|
| PK666 | ITK | Y512 | -16 |
| PN110 | CRYAB | S45 | -17 |
| PN574 | BCLAF1 | S512 | -17 |
| NK030-2 | CDK7 | Pan-specific | -19 |
| PN512 | ENO2 | Y25 | -20 |
| PK558 | CDC7 | T376 | -22 |
| PK793 | Ret (GDNF receptor) | Y905 | -23 |
| PN002 | ACC1 (ACACA) | S80 | -23 |
| NK052-5 | EGFR (ErbB1) | Pan-specific | -26 |
| PN590 | FOXK1 | S441+S445 | -31 |
| PK570 | CDK5 | Y15 | -34 |
| NP001 | CD45 (PTPRC; Receptor-type tyrosine-protein phosphatase C) | Pan-specific | -38 |
| PN196 | GATA1 | S142 | -40 |
| PN144 | PLCG1 | Y783 | -42 |

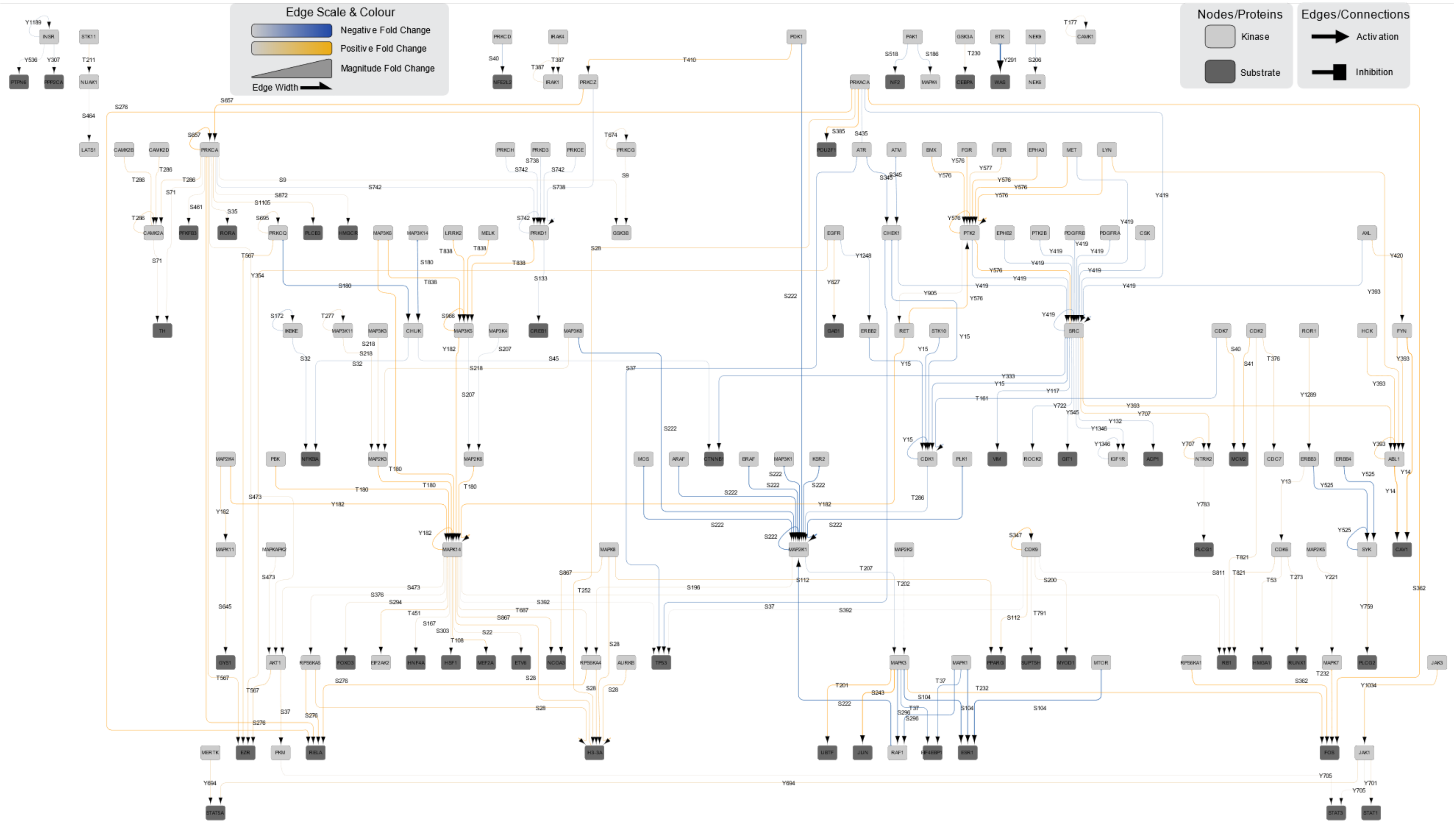


**Figure S3: Microarray map for the A549 sample**

**Table S2: Table listing the main leads for the microarray A549 sample.**

| Antibody ID No. | Target Name | Antibody P-Site | Log2 Fold Change |
|---|---|---|---|
| sc-7263 | Wip1 (PPM1D) | Pan-specific | 2,26 |
| NK116-3 | mTOR (FRAP) | Pan-specific | 2,12 |
| NK225-2 | MEKK6 (MAP3K6; ASK2) | Pan-specific | 1,68 |
| PN827 | Huntingtin (HTT) | Pan+S13+S16 | 1,57 |
| sc-6241 | HSP105 (HSPH1; HSP110) | Pan-specific | 1,52 |
| NK165-3 | RSK2 (RPS6KA3) | Pan-specific | 1,52 |
| sc-639 | PKD1 (PRKCM; PKCm; PRKD1) | Pan-specific | 1,51 |
| PK917 | TrkB (NTRK2) | Y706+Y707 | 1,49 |
| KAP-ST205 | IRAK2 | Pan-specific | 1,42 |
| sc-7973 | p38a MAPK (MAPK14; CSBP; MXI2; SAPK2a) | Y182 | 1,41 |
| NNCOV2S-5 | SARS-CoV-2 Spike RBD | Pan-specific | 1,41 |
| 07-191 | H2B (Histone H2B) | S15 | 1,39 |
| NK229-2 | MELK | Pan-specific | 1,39 |
| PK882 | GSK3a | Y279 | 1,37 |
| PK910 | PKR1 (PRKR; EIF2AK2) | T451 | 1,34 |
| sc-639 | PKD1 (PRKCM; PKCm; PRKD1) | Pan-specific | 1,29 |
| PK893 | GCK (Glucokinase) | Pan+S411 | 1,29 |
| PK537 | GRK2 (BARK1; ADRBK1) | Y356 | 1,28 |
| NK004-3 | CDK15 (PFTAIRE2; ALS2CR7) | Pan-specific | 1,28 |
| PN629 | NCOA3 (SRC-3) | S867 | 1,26 |
| PK571 | CDK6 | Y13 | 1,25 |
| NK105-4 | MKK6 (MAP2K6; MEK6) | Pan-specific | 1,25 |
| NK104-5 | MEK5 (MAP2K5; MKK5) | Pan-specific | 1,24 |

| | | | |
|---|---|---|---|
| PN700 | FBPase 2 (FBP2) | Y216 | 1,23 |
| PN824 | H2AFX (H2AX; Histone H2A.X) | S140 | 1,22 |
| PN644 | PPARg-1 | S112 | 1,21 |
| NN265-1 | Grp170 (HYOU1; ORP-150) | Pan-specific | 1,21 |
| PK647 | GSK3a | S278+Y279 | 1,21 |
| NNCOV2S-9 | SARS-CoV-2 Spike S2 | Pan-specific | 1,20 |
| PN782 | UGDH | Y352 | 1,18 |
| PK712 | Met (HGF receptor) | Y1234+Y1235 +S1236 | 1,16 |
| NK116-5 | mTOR (FRAP) | Pan-specific | 1,16 |
| 44-956 | PKCg (PRKCG) | T514 | 1,15 |
| NK225-3 | MEKK6 (MAP3K6; ASK2) | Pan-specific | 1,12 |
| 05-184 | Src | Pan-specific | 1,12 |
| PK567 | CDK12 (Cdc2L7; CRK7) | T893 | 1,11 |
| NK107-3 | MEKK1 (MAP3K1) | Pan-specific | 1,10 |
| PN772 | PKM2 | Y105 | 1,10 |
| PK574 | CDK9 | S347 | 1,07 |
| PN718 | PKM2 | S37 | 1,07 |
| PN703 | GYS1 | S641+S645 | 1,07 |
| NK025-7 | CDK1 (CDC2) | Pan-specific | 1,06 |
| PK729 | mTOR (FRAP) | S2448 | 1,06 |
| NK237-2 | ATR | Pan-specific | 1,05 |
| NK059-5 | p38g MAPK (MAPK12; ERK6; SAPK3) | Pan-specific | 1,04 |
| PK606 | EphA2 | Y588 | 1,03 |
| NN060-12 | Hsc70 (HSPA8; Hsc70; HSP73; HSPA10) | Pan-specific | 1,03 |
| NK112 | Mos | Pan-specific | 1,03 |
| NK280-1 | MLK4 (MAP3K21) | Pan-specific | 1,03 |
| sc-6212 | Ksr1 | Pan-specific | 1,02 |
| PN719 | PLCB3 | S1105 | 1,02 |
| NK283-1 | MRCKb (CDC42BPB) | Pan-specific | 1,02 |
| AP7642a | VEGFR1 (Flt1) | Pan-specific | 1,01 |

| | | | |
|---|---|---|---|
| NK033-3 | CDK10 (PISSLRE) | Pan-specific | 1,01 |
| NK269-2 | Frk | Pan-specific | 1,01 |
| PK770 | PKD1 (PRKCM; PKCm; PRKD1) | S205 | 1,01 |
| 06-0032 | STAT5A | Y694 | 1,00 |
| NK275-1 | MARK2 | Pan-specific | -1,00 |
| 9111 | CDK1 (CDC2) | Y15 | -1,00 |
| PN637 | TP53 (p53) | S6+S9 | -1,01 |
| PN630 | NFAT1 | S217+S221 | -1,02 |
| PN710 | NMDAR2A NMDA (GRIN2A; Glutamate [NMDA] receptor subunit epsilon-1) | Y943 | -1,02 |
| PN514 | ESYT1 | Y822 | -1,02 |
| PN584 | ERF | T526 | -1,03 |
| PN513 | ERBB2IP (Erbin) | Y1104 | -1,03 |
| NNCOV2S-16 | SARS-CoV-2 Spike S2 | Pan-specific | -1,05 |
| AP7611b | EphA6 | Pan-specific | -1,05 |
| PK624 | ERK4 (MAPK4) | S186 | -1,06 |
| NP008-2 | DUSP2 (PAC1) | Pan-specific | -1,06 |
| PN864 | ARID1A | Y1508 | -1,07 |
| PK792 | Raf1 (c-Raf; RafC) | S301+T303 | -1,07 |
| PN698 | FASN (FAS) | S207 | -1,08 |
| PK638 | Fgr | Y208+Y209 | -1,09 |
| PK747 | p70S6Kb (S6Kb2; RPS6KB2) | S423 | -1,10 |
| PK880 | ERK2 (MAPK1; ERT1) | Y263+S266 | -1,11 |
| NK120-8 | p38a MAPK (MAPK14; CSBP; MXI2; SAPK2a) | Pan-specific | -1,11 |
| sc-7439 | Nek2 | Pan-specific | -1,12 |
| 11097 | TP53 (p53) | S33 | -1,12 |
| PK736 | NLK | T298 | -1,14 |
| 07-012 | FAK (PTK2) | Y397 | -1,15 |
| AP7518b | CDK2 | Pan-specific | -1,16 |
| PK866 | ERK1 (MAPK3; ERT2) | Y204+T207 | -1,16 |
| PN761 | NF1 | Y2577 | -1,16 |
| NK019-3 | CAMK2d | Pan-specific | -1,17 |

| | | | |
|---|---|---|---|
| NK059-4 | p38g MAPK (MAPK12; ERK6; SAPK3) | Pan-specific | -1,18 |
| AAP-104 | CASP4 (Caspase 4) | Pan-specific | -1,18 |
| PN709 | NF2 | S518 | -1,19 |
| NK085-4 | JAK2 | Pan-specific | -1,19 |
| NN604-2 | NRP1 | Pan-specific | -1,21 |
| AP7802b | IRAK1 | Pan-specific | -1,22 |
| NK231 | ErbB3 (HER3) | Pan-specific | -1,22 |
| NN454-1 | 14-3-3-S (YWHAS; SFN) | Pan-specific | -1,22 |
| PN631 | NFAT3 (NFATc4) | S213+S217 | -1,23 |
| DB033 | NFKB p65 (Rel A) | Pan-specific | -1,23 |
| PN841 | NMDAR1 (NR1) | S897 | -1,28 |
| PK743 | p38d MAPK (MAPK13) | Y182 | -1,28 |
| NN300-1 | NrCAM | Pan-specific | -1,28 |
| 11134 | CDK1 (CDC2) | T161 | -1,28 |
| AP7612a | EphA7 | Pan-specific | -1,30 |
| PK873 | Abl (Abl1) | Y393+T394 | -1,30 |
| 11063 | EZR (Ezrin; VIL2) | Y354 | -1,30 |
| PK629 | FAK (PTK2) | Y577 | -1,31 |
| NK120-10 | p38a MAPK (MAPK14; CSBP; MXI2; SAPK2a) | Pan-specific | -1,33 |
| C27220 610266 | CD45 (PTPRC; Receptor-type tyrosine-protein phosphatase C) | Pan-specific | -1,33 |
| 11098 | TP53 (p53) | S37 | -1,35 |
| sc-7230 | ATM | Pan-specific | -1,35 |
| PK879 | ERK1 (MAPK3; ERT2) | S283 | -1,36 |
| sc-1214 | ATM | Pan-specific | -1,37 |
| NN296-1 | NLGN4x (NLGN4l) | Pan-specific | -1,37 |
| NK121-4 | p38d MAPK (MAPK13) | Pan-specific | -1,39 |
| PN842 | NRF2 | S40 | -1,42 |
| AP7500a | ERK1 (MAPK3; ERT2) | Pan-specific | -1,44 |
| PK627 | FAK (PTK2) | Y397 | -1,45 |
| PK886 | MEK1 (MAP2K1; MKK1) | T286 | -1,46 |
| NN229-1 | CD74 | Pan-specific | -1,50 |
| PP527 | CD45 (PTPRC; Receptor-type tyrosine-protein phosphatase C) | Y1216 | -1,50 |

| | | | |
|---|---|---|---|
| NN451-3 | 14-3-3e (YWHAE) | Pan-specific | -1,52 |
| AP7504a | ERK5 (MAPK7; BMK) | Pan-specific | -1,54 |
| PK865 | ERK1 (MAPK3; ERT2) | T207 | -1,56 |
| NN230-1 | CDC37 | Pan-specific | -1,61 |
| PK878 | ERK1 (MAPK3; ERT2) | S265 | -1,62 |
| PK558 | CDC7 | T376 | -1,65 |
| NP038-1 | CDC25A | Pan-specific | -1,66 |
| 44-864 | Integrin a4 - pS1021 | S1021 | -1,68 |
| NP038-3 | CDC25A | Pan-specific | -1,76 |
| PK888 | p38d MAPK (MAPK13) | S261+T265 | -1,80 |
| DB040 | Trail | Pan-specific | -1,81 |
| C25820 610250 | CDC34 | Pan-specific | -1,88 |
| IMG-139 | TBK1 (IKKd) | Pan-specific | -1,88 |
| NN204-1 | ATAD1 ATPase | Pan-specific | -1,94 |
| PK791 | Raf1 (c-Raf; RafC) | S296 | -2,02 |
| PN521 | ITSN2 | Y968 | -2,03 |
| PK668 | JAK2 | Y570 | -2,07 |
| NK059-3 | p38g MAPK (MAPK12; ERK6; SAPK3) | Pan-specific | -2,15 |
| NK020-1 | CaMK2g | Pan-specific | -2,18 |
| sc-1284 | ERK5 (MAPK7; BMK) | Pan-specific | -2,19 |
| NN205-2 | Ataxin 1 (Atxn1; SCA1) | Pan-specific | -2,19 |
| PK555 | CaMK2a (CaMKII) | T286 | -2,28 |
| NP038-2 | CDC25A | Pan-specific | -2,29 |
| 13-9800 | CAMK2b | Pan-specific | -2,32 |
| DB075 | IkBa (MAD3; IkBa) | Pan-specific | -2,49 |
| sc-58758 | EZR (Ezrin; VIL2) | Pan-specific | -2,52 |
| sc-1285 | ERK5 (MAPK7; BMK) | Pan-specific | -2,54 |
| NK055-2 | ERK1 (MAPK3; ERT2) | Pan-specific | -2,60 |
| sc-56070 | CASP8 (Caspase 8) | Pan-specific | -2,75 |
| 13-9800 | CAMK2b | Pan-specific | -2,95 |
| NK084-3 | JAK1 | Pan-specific | -3,00 |

| | | | |
|---|---|---|---|
| sc-56063 | CASP7 p20 (Caspase 7) | Pan-specific | -3,03 |
| PK659 | IKKa (IkBKA) | T179+S180 | -3,55 |
| sc-622 | CASP1 (Caspase-1) | Pan-specific | -4,17 |

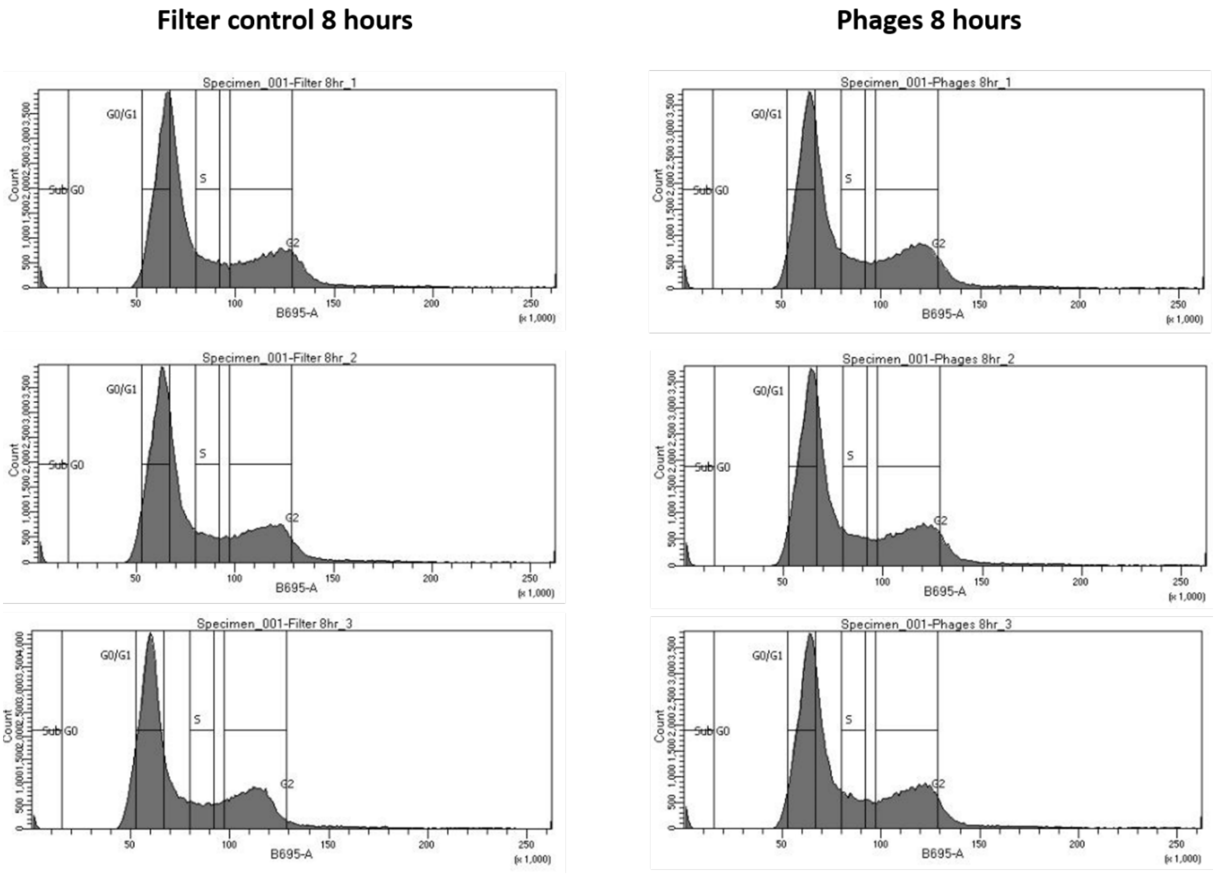


**Figure S4: FACS count results showing the distribution of the cells across the different cell cycle stages.** On the left, the three control samples were incubated with the Filter control for 8 hours. On the right, the three samples were incubated with the T4 phages for 8 hours. The cell cycle stages were set on a non-incubated sample and kept fixed for all the following analyses.

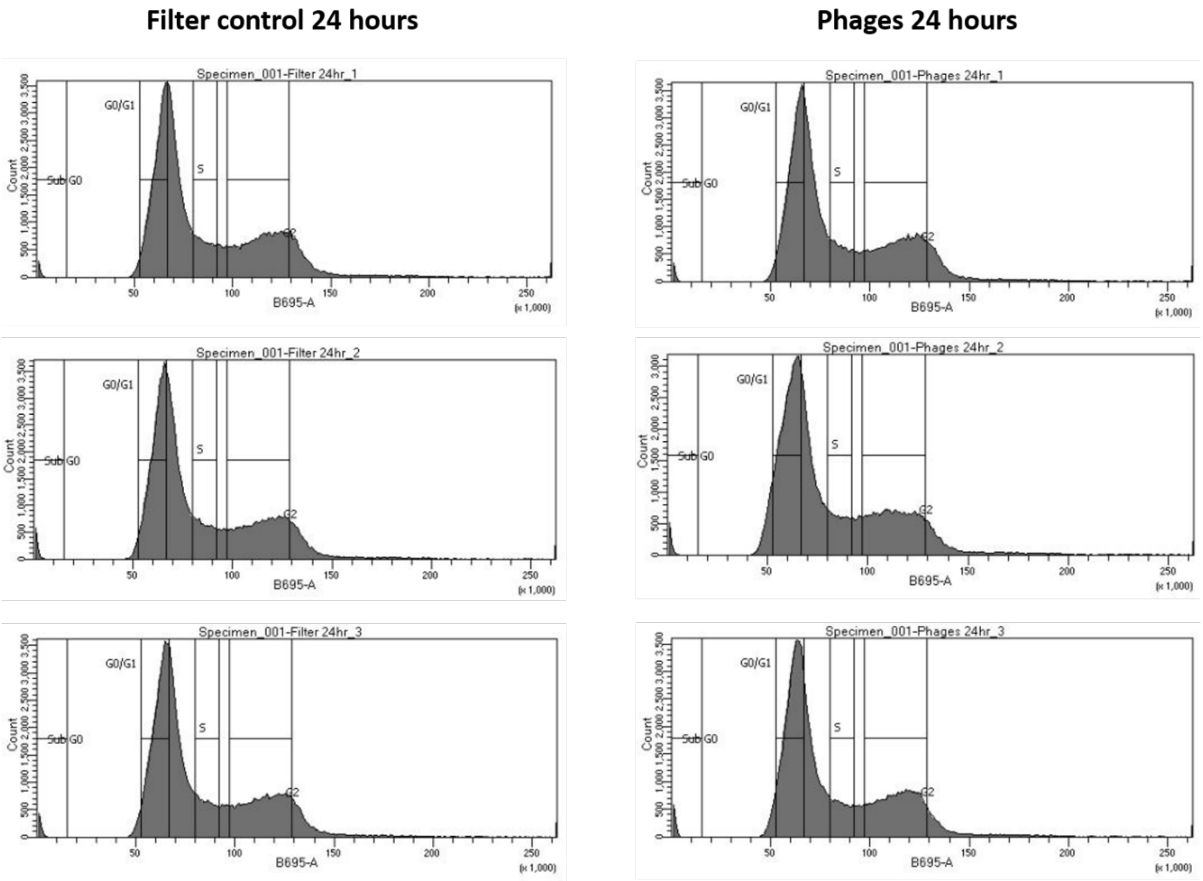


**Figure S5: FACS count results showing the distribution of the cells across the different cell cycle stages.** On the left, the three control samples were incubated with the Filter control for 24 hours. On the right, the three samples were incubated with the T4 phages for 24 hours. The cell cycle stages were set on a non-incubated sample and kept fixed for all the following analyses.
